# Investigating fine-scale breeding habitat use by amphibians in a continuous wetland using environmental DNA

**DOI:** 10.1101/2023.02.27.530201

**Authors:** Julie Morgane Guenat, Antoine Gander, Luca Fumagalli, Guillaume Lavanchy

## Abstract

Designing effective conservation plans to protect species from extinction requires a better understanding of their ecology. Conventional methods used to investigate habitat use are time consuming, and detectability of cryptic species is often insufficient. Environmental DNA (eDNA)-based approaches now provide an alternative for ecosystems monitoring and assessment. Nevertheless, to our knowledge, such methods have never been applied to investigate habitat use at a fine scale in a continuous wetland environment. Here, we used an eDNA metabarcoding approach to characterize the breeding habitat use of local amphibian species in a wet meadow expanse along the southern shore of Lake Neuchâtel, Switzerland. We retrieved DNA from six out of the seven species expected to be present. We tested the influence of six abiotic environmental variables on overall species communities as well as individual species occurrences. We showed that the main factor structuring species communities was water temperature, and that the distribution of three amphibian species was associated with several environmental variables. Our results indicate that the eDNA approach is a promising tool to study species’ ecology at a small scale in continuous wetland habitats.

## 1 INTRODUCTION

Amphibian populations have been collapsing for decades (Houlahan et al., 2000; Zedan, 2004). This decline is partly attributed to habitat loss and degradation (Arntzen et al., 2017; Becker et al., 2010). While most amphibians live in freshwater wetlands or moist habitats, 87% of global wetland surface has been destroyed during the last three centuries (Davidson, 2014). Conservation measures to preserve the remaining natural habitats are therefore crucial to reverse current population declines.

Designing sound conservation measures requires a deep understanding of species ecology and population trends (Cristescu & Boyce, 2013). Thus, reliable and efficient monitoring methods are needed. However, standard amphibian surveys relying on acoustic and/or visual detection and trapping are time consuming, and often prove to be inefficient to monitor cryptic species, such as most Urodela species (Rödel & Ernst, 2004). Hence, effective survey tools must be developed to increase efficiency and detectability of such species.

In this perspective, recently developed environmental DNA (eDNA)-based survey methods offer new tools for monitoring wildlife communities (Garlapati et al., 2019; Ruppert et al., 2019; Taberlet et al., 2018). The eDNA approach is a non-invasive survey method that relies on the collection and analysis of DNA released by individuals through dead cells, hair and feces into the environment (e.g., soil or water; Taberlet et al., 2012). This approach, combined with DNA metabarcoding (i.e. the amplification and sequencing of short species-diagnostic DNA sequences with universal primers; Taberlet et al., 2012) has been used for current and ancient biodiversity inventories (e.g., Lodge et al., 2012; Willerslev et al., 2003), to survey endangered (e.g., Thomsen et al., 2012) and invasive species (e.g., Dejean et al., 2012; Ficetola et al., 2008; Smart et al., 2015), in diet analyses (e.g., De Barba et al., 2014; Shehzad et al., 2012), but also to study trophic relationships and niche partitioning (e.g., Calderón-Sanou et al., 2021; Chalmandrier et al., 2019; Martinez-Almoyna et al., 2019) and habitat use (e.g., Perl et al., 2022; Sakata et al., 2017).

eDNA-based methods are generally more sensitive and efficient, and often more cost-effective, than standard monitoring methods (Biggs et al., 2015; Lopes et al., 2017) and have been largely used to monitor amphibian species (reviewed in Ficetola et al., 2019). Nevertheless, for these taxa, eDNA metabarcoding approaches have so far only been used in discrete environments such as distinct water bodies (e.g., Biggs et al., 2015; Dufresnes et al., 2019; Eiler et al., 2018), or streams (e.g., Li, et al., 2021a; Pilliod et al., 2013, 2014) to monitor species occurrences, but never in continuous natural wetlands to study habitat use at a fine scale.

In this study, we used eDNA metabarcoding to study the fine scale breeding habitat use of the amphibian species present in a continuous wetland expanse in Switzerland. We tested the influence of six abiotic environmental variables (average mud depth, percentage of emerged and submerged vegetation, percentage of emerged land, average water temperature and the distance to the wintering habitat) on community structure and individual species distribution. Finally, we investigated whether biotic interactions could influence amphibian distribution by examining co-occurrences of the different species in the studied area.

## 2 METHODS

### 2.1 Study area

*La Grande Cariçaie*, located along the southern shore of Lake Neuchâtel, is the largest remaining continuous wetland area in Switzerland. The area is protected by eight natural reserves, totaling about 660 ha. The habitat consists of three main vegetation types (after Delarze et al., 1998): sedge meadows dominated by *Carex elata* and *Cladium mariscus* (*Magnocaricion*); reedbeds dominated by *Phragmites australis* (*Phragmition*); and open water bodies (ponds, ruts) dominated by *Nymphaea alba* (*Nymphaeion*). In the present study, we focused on two out of the eight natural reserves: Yverdon and Gletterens (Fig. 1).

**Figure 1:**
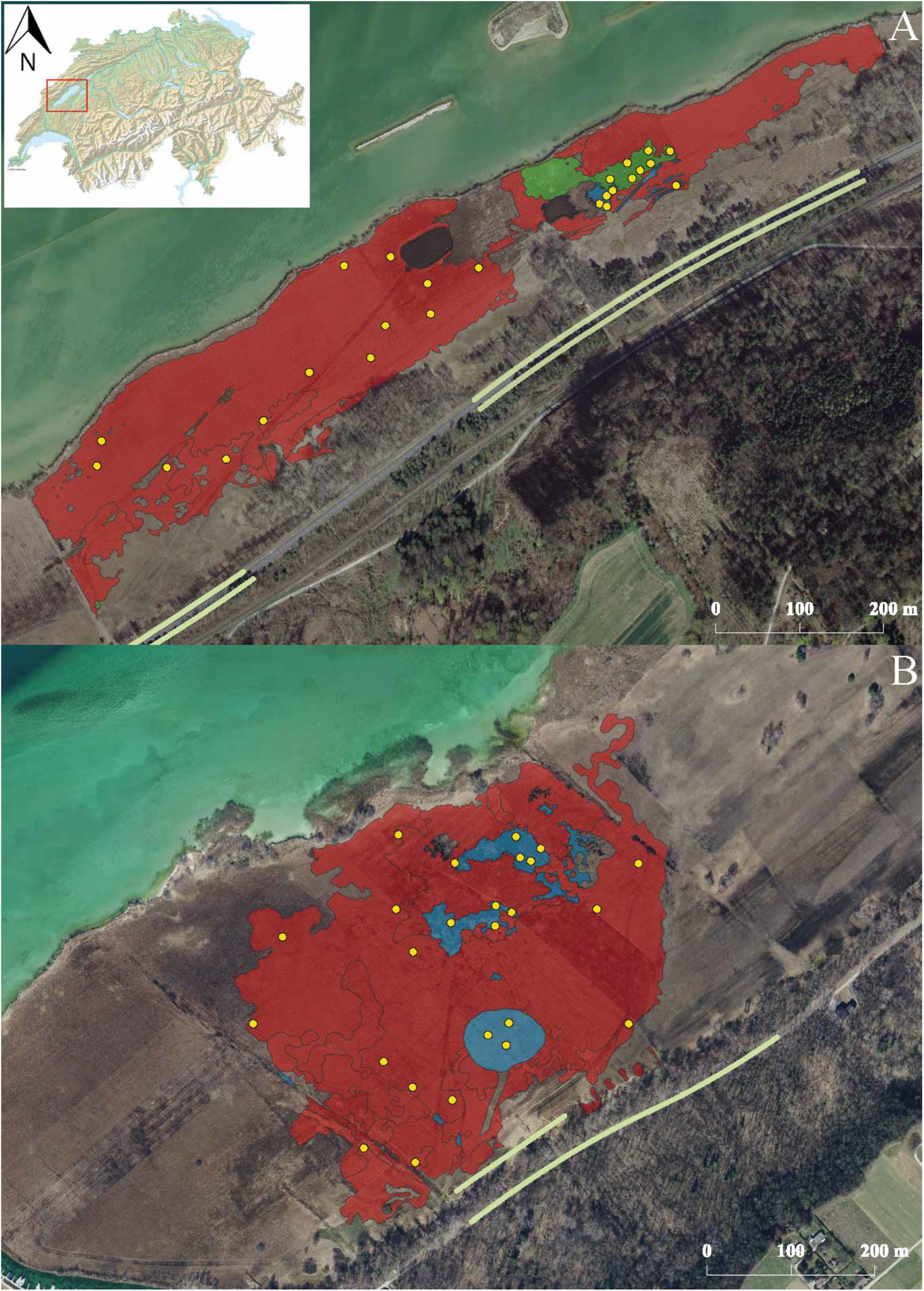
Location of the study areas. Top left: map of Switzerland with the study area delineated by a red square. **A** Map of Yverdon. **B** Map of Gletterens. Habitat types (sensu Delarze and Gonseth 1998) are shown by coloured polygons (red: *Magnocaricion*; green: *Phragmition*; blue: *Nymphaeion*). Yellow dots represent sampling points. Green lines represent amphibian barriers used for the prenuptial migration study. The background map was extracted from Google map using QGIS.

Seven amphibian species are present in the *Grande Cariçaie*: *Lissotriton vulgaris, L. helveticus, Bufo bufo, Rana temporaria, Hyla arborea* and the *Pelophylax* complex comprising two mitochondrial haplotypes: *P. ridibundus* and *P. bergeri* (Dubey et al., 2014; Dufresnes et al., 2017).

To confirm the presence and distribution of amphibian species along the study area, as well as to obtain an estimate of their densities, we monitored the number of individuals per species during prenuptial migration using interception barriers (Table S1). Local amphibian species, except most *Pelophylax*, winter in forests and migrate in wetlands to breed. Wintering habitats in Yverdon are separated from the breeding habitat by a road. To reduce mortality due to traffic, barriers leading to tunnels allowing amphibians to cross below the road were built. These tunnels are on average 58.28 m apart (min = 35 m; max = 105 m) over 408 m, i.e., totaling seven tunnels. We placed net traps at the exit of the tunnels to capture and count migrating individuals every morning from January 23^rd^ to April 4^th^, 2018 (with some interruptions when migration was stopped by low temperatures). In Gletterens, we installed a 420 m long barrier between the forest and the wetlands to intercept migrating individuals, which then fell into buckets buried every approx. 25 m. We then counted them and released them beyond the barrier. For logistic reasons, the barrier was set only between March 1^st^ and March 31^st^, 2018, which can result in an underestimation of the number of individuals during the prenuptial migration compared to the other location. However, it still provides a reliable estimation of the relative density of amphibians along the sampled area.

We defined a sampling point as a 5 m diameter circle within which we collected water samples and measured the environmental variables that were used to characterize the habitat. At both locations, we randomly assigned 25 sampling points stratified across the three vegetation types (*Magnocaricion, Phragmition* and *Nymphaeion*; Fig. 1).

These are referenced in a vegetation map produced between 2012 and 2014 using a combination of aerial photographs and field identification of the different habitat types (based on (Delarze et al., 1998). To ensure independence of our samples, the centers of all sampling points were located at least 20 m apart. Due to the relatively small surfaces of *Phragmition* and *Nymphaeion*, fewer samples could be collected in these habitats than in *Magnocaricion* (final number of sampling points: 27 for *Magnocaricion*, 16 for *Nymphaeion* and 7 for *Phragmition* (Fig. 1)).

### 2.2 Water sampling

We collected water samples from May 21^st^ to 28^th^ 2018, which corresponds to the breeding season for local amphibians. We collected two liters of water at each sampling point using the VigiDNA kit (Spygen, Le Bourget du Lac, France) following manufacturer’s protocol. We were interested in recent amphibian presence (i.e., individuals present during the breeding season). In the water column, DNA becomes undetectable within approximately two weeks (Dejean et al., 2011; Thomsen et al., 2012), but it can be preserved for much longer in the sediments (Barnes & Turner, 2016; Nielsen et al., 2007). To avoid resuspending sediments while walking near the sampling points, we attached the spoon to collect water provided by the kit to a 4 m fishing rod (Fig. S1). To avoid cross-contamination between sampling points, we ensured the fishing rod did not enter the water and rinsed it with bottled water after sampling. Filtration capsules were stored at room temperature for two months, until DNA extraction.

### 2.3 Environmental variables

To characterize the habitat at each sampling point, we measured five environmental variables: average mud depth, percentage of emerged and submerged vegetation, percentage of emerged land and average water temperature. In addition, we computed the distance to the wintering habitat (nearest forest) using QGIS (version 3.0.1).

We estimated average mud depth by averaging measures taken at the center and at 2.5 m from the center at the four cardinal points and at the four edges of the sampling point (Fig. S2). The same observer estimated coverage of emerged and submerged vegetation as well as emerged land for all points. We measured these environmental variables from April 21^st^ to 23^rd^ 2018, i.e., one month prior to water sample collection to avoid contamination with amphibian DNA from sediments released during this measurement procedure. We measured the water temperature every hour at each point simultaneously during the sampling period (April 21 to May 28), which also corresponds to the time of egg laying and development of amphibian larvae (Meyer et al., 2009). At each sampling point, we placed thermologgers (1-Wire®/iButton) in Falcon tubes and sealed them with parafilm (Supplementary Methods S1). We then calculated the average temperature for each sample point over the entire measurement period.

### 2.4 Laboratory procedures

We extracted the DNA from the VigiDNA capsules in a room dedicated to low DNA-content samples extraction and pre-PCR setup. We followed the DNA extraction protocol as described in (Mas-Carrió et al., 2022). We eluted DNA by adding 100 μL of SE buffer and centrifuging twice.

Before performing the amplification of the extracted DNA, we tested eight samples chosen randomly among the 50 samples for inhibitors using RT-qPCR (Biggs et al., 2015), carried on a QuantStudio 3 Real-Time PCR instrument (ThermoFisher). We amplified a fragment of the 12S mitochondrial ribosomal DNA using the BATR01 primers (Valentini et al., 2016). The qPCR mixture contained 1x AmpliTaq ^™^ Gold 360 mix (Applied Biosystem ^™^), 0.5 μM of BATR01 primers and 10,000-times diluted stock SyberGreen (ThermoFischer Scientific). Since BATR01 primers amplify other vertebrate species including *Homo sapiens*, we added to the qPCR mix 2 μM of human-blocking primer (batra_blk; Valentini et al., 2016). The final volume was 20 μL including 2 μL of DNA template. We diluted the eight samples 1x, 0.5x or 0.1x and replicated each concentration three times. We included four PCR and four extraction negative controls in the qPCR plate as well as a dilution series (from 1x to 10^−5^x of *R. iberica* DNA extracted from tissues with an initial concentration of 47.2 ng/μL) replicated three times. Thermocycling conditions were the following: denaturation at 95°C for 10 min, followed by 55 cycles of 30 s at 95°C, 30 s at 55°C and 1 min at 72°C, followed by a melting curve analysis of the PCR products by heating from 55°C to 95°C and continuous measurement of the fluorescence. The samples showed no inhibition, hence, we did not dilute the 50 samples for further metabarcoding amplifications.

We then amplified the extracted DNA from the 50 water samples. The PCR mix included 1x AmpliTaq^™^ Gold 360 mix (Applied Biosystem^™^), 0.5 μM of each dual combinatorial indexing tagged forward and reverse BATR01 primers (i.e., primers with eight variable nucleotides added to their 5’ end, allowing further sample identification) and 2 μM of human-blocking primer. The final volume was 20 μL including 2 μL of DNA template. We replicated each sample amplification 12 times in 12 separate PCR plates. In each PCR plate, we set 12 blanks corresponding to empty wells and allowed to estimate the proportion of sequence leaking and tag switches (i.e., false combination of tags, generating chimeric sequences occurring during the sequencing process), seven negative extraction controls (extraction performed on Milli-Q^(R)^ water), seven negative PCR controls (water) and seven positive controls containing an equimolar assembly of DNA from three exotic species (*P. nigromaculatus, Pseudacris maculata* and *R. arvalis*), with a similar DNA concentration to that of eDNA samples estimated with the qPCR (Supplementary methods S2; see Taberlet et al., 2018 for plate layout). Thermocycling conditions were: denaturation at 95°C for 10 min, followed by 40 cycles of 30 s at 95°C, 30 s at 55°C and 1 min at 72°C, with a final elongation step of 7 min at 72°C. One out of the seven positive and negative PCR controls per replicate plate was visualized on a 1.5 % agarose gel stained by ethidium bromide. We excluded one of the twelve replicate PCR plates from further analyses since no amplification was detected. We then pooled the PCR products from the eleven remaining plates, and purified amplicons using a MinElute PCR purification kit (Qiagen).

We size-selected BATR01 amplicons on a 2% agarose gel and purified them using MinElute Gel Extraction kit (Qiagen). Library preparation was performed using TruSeq® DNA PCR-Free Library Prep (Illumina) with the following modifications to ensure a maximal yield of small-size DNA (amplicon size is on average 110 bp including primers): (i) we skipped the “Remove large fragments” step; (ii) we added 100 μL of undiluted SPB to the 100 μL of end-repaired sample; (iii) and we followed the protocol starting from step three of the “Remove small fragments” step. The final library was quantified by qPCR using the KAPA Library Quantification Kit (Roche). Sequencing of the library was carried out at the Genomic Technologies Facility (University of Lausanne, Switzerland). A 100 cycles paired-end sequencing was performed on a single Illumina HiSeq 2500 lane.

### 2.5 Sequence processing

We compiled a reference database by recovering the set of vertebrate DNA sequences from EMBL-European Nucleotide Archive (release 138) and by downloading Taxonomy from NCBI. We converted those files into ecoPCR format using *obiconvert* (OBITools; Boyer et al., 2016) and performed an *in-silico* PCR using ecoPCR (Gentile F. Ficetola et al., 2010). We retained all sequences that (i) matched BATR01 primers with up to three mismatches, except in the two last nucleotides of the primer, and (ii) had a minimum and a maximum amplicon length between 15 bp and 101 bp, respectively (Bellemain et al., 2010; Valentini et al., 2016).

The metabarcode sequence used in the present study was missing for *L. helveticus* and for *P. bergerii* in the reference database. Thus, we Sanger-sequenced a fragment of the mitochondrial 12S gene covering the whole length of our metabarcode using DNA extracted from a tissue sample for both species and amplified using primers L2519 and H3296 (Wang et al., 2017)(accession number: *L. vulgaris* OK032503 and *P*. bergerii OQ508898). We then added the sequences manually to the reference database.

Sequence reads were processed using OBITools (Boyer et al., 2016). We assembled forward and reverse reads using *illuminapairedend* with a minimal quality score set to 40 and discarded unaligned sequences (i.e., labeled as joined sequences) using *obigrep*. We assigned each sequence to samples (i.e., we demultiplexed them) using *ngsfilter*. We dereplicated reads by clustering strictly identical sequences into a unique sequence using *obiuniq*. We removed singletons and assigned molecular operational taxonomic units (MOTUs) to a taxon using *ecotag* with the reference database. We cleaned the taxonomically assigned sequences from PCR and sequencing errors using *obiclean* with a minimum ratio between counts of two sequence records set at 0.25.

All downstream analyses were conducted in R version 4.0.3 (R core Team & R Foundation for Statistical Computing, 2020). We first removed unassigned sequences and sequences with an identity lower than 98%. We then used several approaches to filter contaminations and chimeric sequences resulting from sequence leaking. To account for contaminations stemming from laboratory manipulation, we retrieved the maximum number of reads across all negative controls and used it as a minimum threshold for presence in a sample. That is, a MOTU was considered present in a sample only if the number of reads was higher than the highest number of reads across all negative controls. Since we retrieved reads corresponding to two exotic species (*Xenopus tropicalis* and *Rhinella sp*.) after these cleaning steps, we removed the number of reads corresponding to these species to all other MOTUs in each sample. We then accounted for sequence leaking by computing and plotting the effect of different proportions of leaking removal on the numbers of MOTUs and sequences (Fig. S3). Based on these graphs we concluded that leaking accounted for 60% of sequences retrieved in samples. Hence, we removed 60% of total reads per MOTUs in each sample.

To consider a species as present, no consensus threshold is set in the literature (Goldberg et al., 2016; Harper et al., 2018). In the present study we attempted to be conservative to limit the occurrence of false positives. We considered a species present if at least 3 out of the 11 PCR replicates contained a positive number of reads after all cleaning steps for a given MOTU.

### 2.6 Statistical analyses

We first asked whether the species were randomly distributed or whether they were aggregated in the study area. We tested for spatial autocorrelation of species assemblies using a Mantel test between a geographic distance matrix and a species assemblage dissimilarity matrix. We used the *vegdist* function from the ‘vegan’ package (Oksanen et al., 2020) to compute pairwise Euclidian distances between the sampling points. We added 0.001 to each cell of the species matrix since five sampling points did not have any amphibian species. We computed the dissimilarity matrix of the species assemblage with the *jaccard* method using *vegdist* function. We performed the Mantel test using the *mantel* function from the package ‘vegan’.

Then, we investigated if the species assemblies could be explained by habitat characteristics. We performed a distance-based redundancy analysis (db-RDA; Legendre & Andersson, 1999), which detects linear relationships between a matrix of explanatory variables and a dissimilarity matrix of response variables generated by measures that are non-linear. We used the function *capscale* from the package ‘vegan’ to perform the db-RDA (Oksanen et al., 2020). In the same way as for the mantel test, we added 0.001 to each species value prior to the analysis. We used the *jaccard* method to compute dissimilarity matrices, as this was found to be the most suitable according to the *rankindex* method from the package ‘vegan’. We tested whether the influence of the environmental variables and their quadratic effects on the species assemblage were significant by performing a permanova (i.e., permutational multivariate analysis of variance) using the function *adonis2* from the package ‘vegan’.

Finally, we aimed at characterizing the breeding habitat of each amphibian species. To this end, we tested the effect of the recorded environmental variables and the effect of other amphibian species on the probability of presence of each amphibian species. We performed separate binomial generalized linear models (GLMs) for each species, using the presence-absence of the focal species as response variable and the environmental variables. The full model contained each environmental variable. To test the effect of other amphibian species on the probability of the focal species, we performed a PCA on the distribution of all other species and used the first axis as an additional explanatory variable. Backwards model selection was then performed using *drop1* from the package ‘lme4’ (Bates et al., 2015), and the model with the lowest AIC was selected. Zero-inflation and GLM assumptions were checked for each model using ‘DHARMa R’ package (version 0.3.3.0; Hartig, 2020).

## 3 RESULTS

After sequencing the DNA extracted from water samples, we obtained 182,672,348 raw reads. After processing them using OBITools, we retained a total of 131,883,594 reads, of which 50,015,762 were attributed to Amphibia, corresponding to a relative read abundance (i.e., proportion of reads, hereafter RRA) of 0.38 which represent the highest RRA among the identified taxa. We identified three other vertebrate classes: Mammalia (RRA = 0.27, of which 99.99% were attributed to *Homo sapiens*), Actinopterygii (RRA = 0.27) and Aves (RRA = 0.02). A small proportion (RRA = 0.06) of sequences were not assigned using our reference database (Fig. 2A).

**Figure 2:**
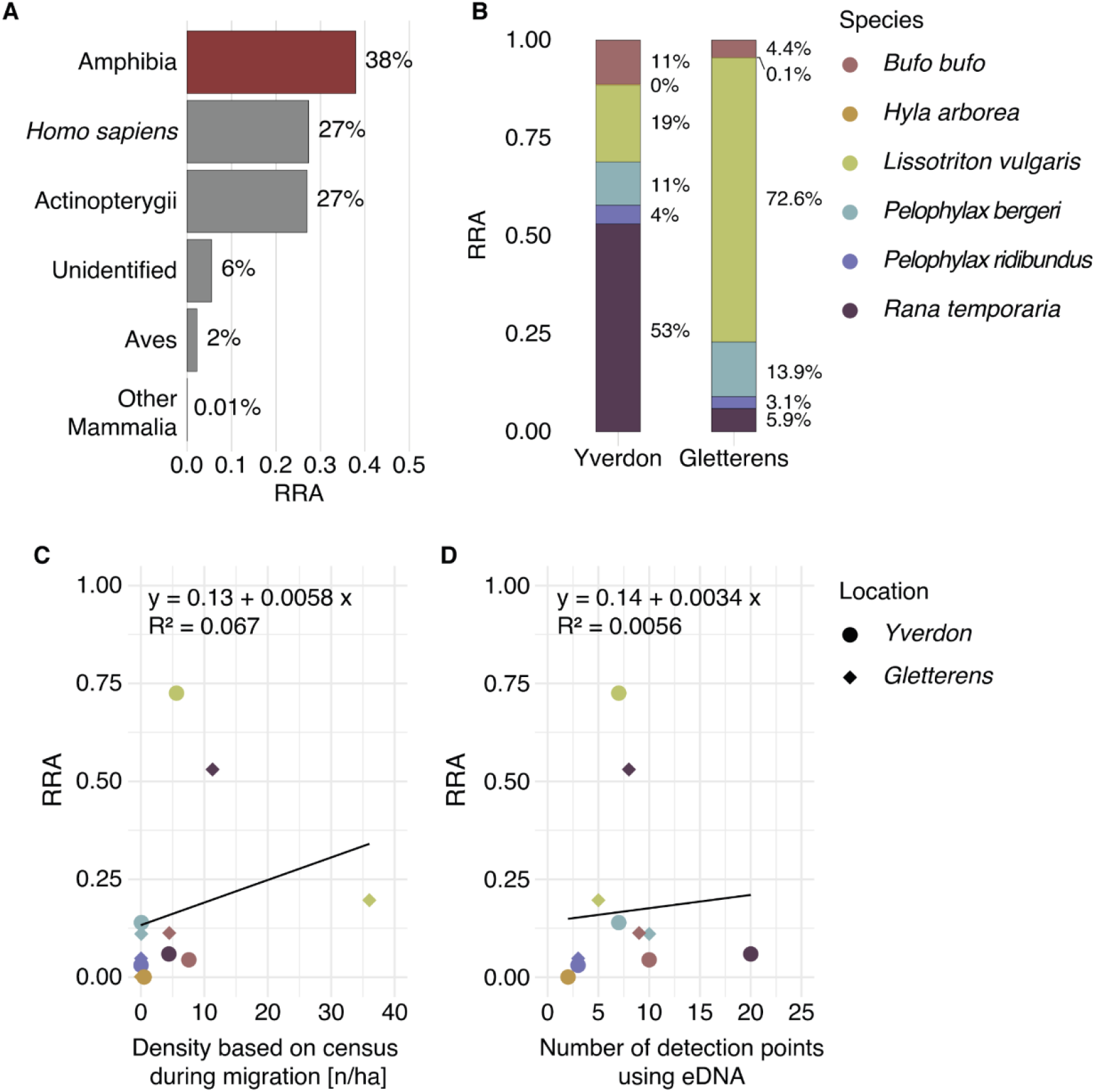
Barplots representing the relative read abundance. **(RRA)** of (**A**) each retrieved class and (**B**) of amphibian species for each of the two locations. (**C**) relation between the RRA of local amphibian species and their estimated densities calculated using the prenuptial migration survey. (**D**) Relation between the RRA of local ampibian species and the number of points where they were detected using eDNA approach. Total number of reads after processing sequences with OBITools software was 182,672,348. The total number of Amphibia reads was 50,015,762.

We detected DNA from six out of the seven species expected in *La Grande Cariçaie* (*H. arborea, B. bufo, L. vulgaris, R. temporaria, P. ridibundus* and *P. bergeri*). No DNA from *L. helveticus* was recovered despite 97 individuals entering the area during the prenuptial migration. In contrast, we detected DNA of *H. arborea* in Yverdon, although no individuals were caught during the prenuptial migration survey (Fig. 2B and Table S1).

Then, we asked whether the density of individuals could be estimated by the amount of DNA sequences recovered for a particular species. We found no correlation between the RRA of each amphibian species and its estimated density calculated from the prenuptial migration survey (*R*^*2*^ = 0.067, *p* = 0.42; Fig. 2C) nor the number of sampling points where the species was detected (*R*^*2*^ = 0.0056, *p* = 0.82; Fig. 2D).

We then investigated which factors explained the distribution of our target species within the wetlands. We found a positive correlation between geographic distance and species dissimilarity between the sampling points (Mantel test; *r* = 0.11, *p* = 0.003). To test whether this correlation could be driven by environmental similarity between nearby plots, we tested whether the species communities were affected by the environmental variables. The db-RDA analysis showed that 27.4% of the variance in species composition could be explained by the recorded environmental variables (Total Inertia Eigenvalue λ_Total_ = 16.67; Constrained Inertia Eigenvalue λ_Constrained_ = 4.57; adjusted *R*^*2*^ = 0.057). The first axis (CAP1, Fig.3) explained 12.77% (λ_1_ = 2.13), the second axis (CAP2, Fig.3) explained 8.04% (λ_2_ = 1.34) and the six remaining constrained axes explained 6.59% of the total variance. The PERMANOVA performed on the db-RDA indicated that the quadratic effect of water temperature seemed to affect the species communities (PERMANOVA: *R*^*2*^ = 0.054, *p* = 0.045, *F* = 2.76; Table S2). The proportion of emerged vegetation as well as the quadratic effect of the proportion of emerged land tended to affect the species assemblies, but these tendencies were not significant (PERMANOVA: *R*^*2*^ = 0.056, *p* = 0.06, *F* = 2.81, and *R*^*2*^ = 0.046, *p* = 0.084, *F* = 2.35, respectively; Table S2).

**Figure 3:**
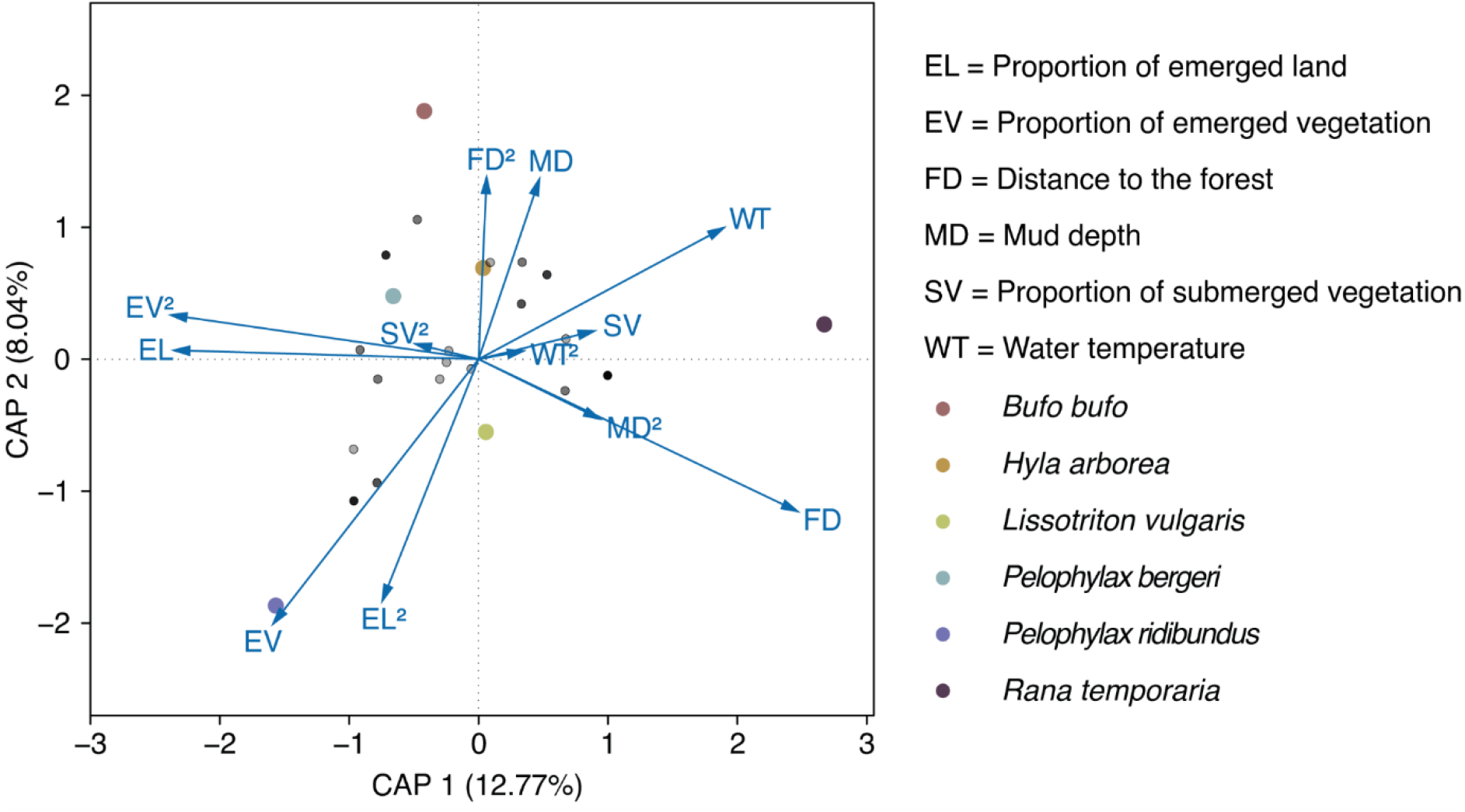
Ordination plot of the distance based-Redundancy Analysis (db-RDA) on the amphibian species composition. Coloured dots represent each species, small transparent grey dots represent the 50 sampling points, blue arrows represent the environmental variables (“^2^” designates the quadratic effect of a variable). 27.4% of the variance in species composition could be explained by the recorded environmental variables (Total Inertia Eigenvalue λ_Total_ = 16.671 and Constrained Inertia Eigenvalue λ_Constrained_= 4.568). The first axis (CAP1) explained 12.77% of the total variance (λ_1_= 2.13), the second axis (CAP2) explained 8.04% (λ_2_= 1.34) and the six constrained axes explained 6.59% of the total variance.

We then investigated the breeding habitat use of each of the local species. The distribution of three out of the six detected species was correlated with some of the environmental variables we recorded (Fig. 4). The distribution of *P. bergeri* was positively affected by the proportion of emerged vegetation (GLM: df = 49, *Z* = 2.052, *p* = 0.04) and tended to be affected negatively by the presence of other anuran species, proxied in the model by the first axis of the PCA on other amphibian species represented in Fig. S4 (GLM: df = 49, *Z* = −1.663, *p* = 0.096). The distribution of *P. ridibundus* was positively associated with high mud depth (GLM: df = 49, *Z* = 2.059, *p* = 0.04). Finally, the distribution of *R. temporaria* was positively affected by the distance to the wintering habitats (GLM: df = 49, *Z* = 2.24, *p* = 0.025). On the other hand, the recorded environmental variables did not seem to affect the distribution of *B. bufo, L. vulgaris* and *H. arborea*, though the distributions of *L. vulgaris* and *H. arborea* tended to be negatively affected by the distance to the wintering habitat (GLM: df = 49, *Z* = −1.79, *p* = 0.074) and vegetation cover (GLM: df = 49, *Z =* −1.15, *p =* 0.071), respectively.

**Figure 4:**
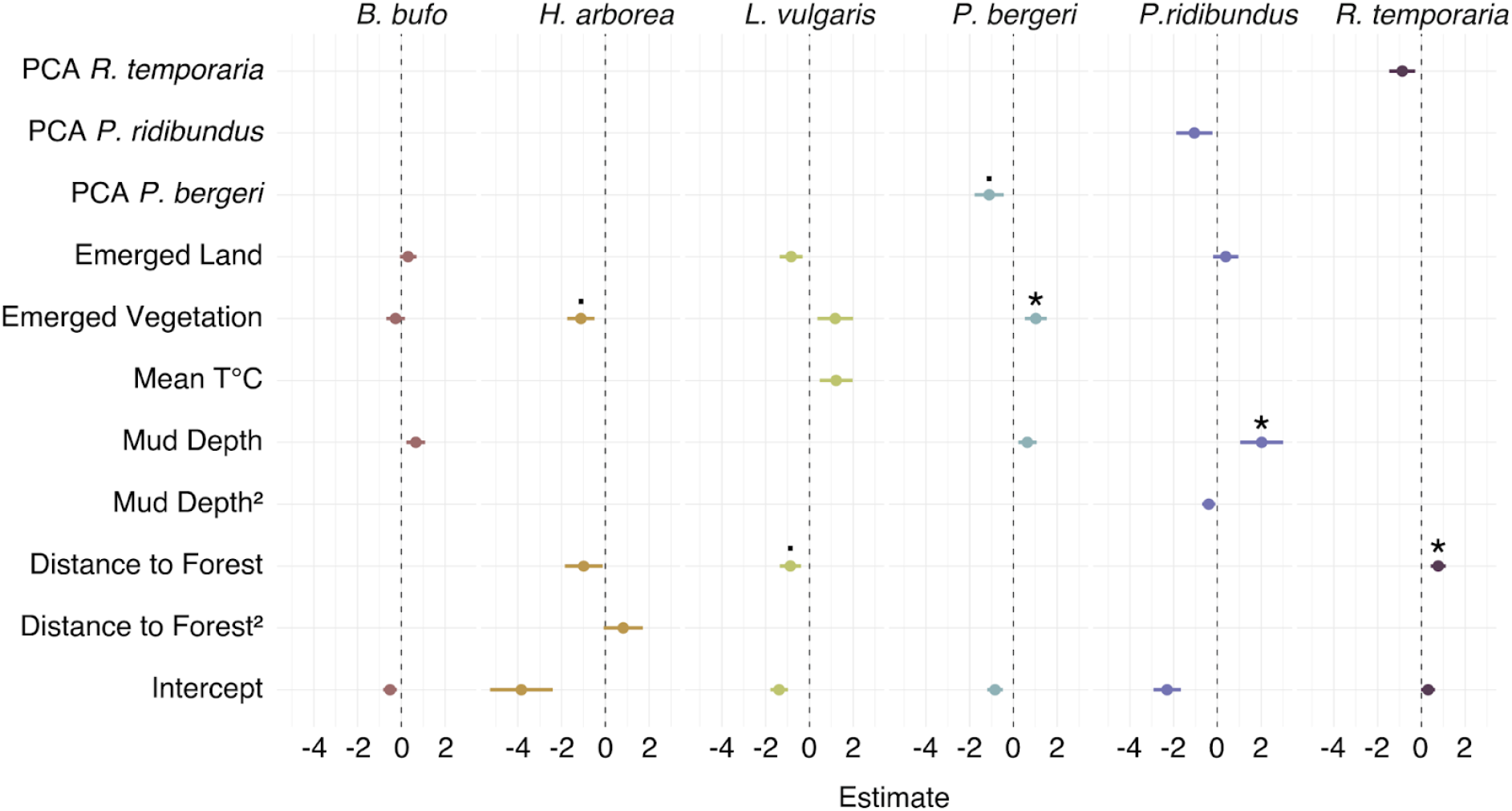
Coefficient plot of the Generalized Linear Models testing the effect of recorded environmental variables on the probability of presence of each of the amphibian species detected using environmental DNA approach. The dots represent the estimate and lines correspond to the standard error. PCA R. temporaria, PCA P. ridibundus and PCA P. bergeri: the effect of other amphibian species on the probability of the focal species, represented by the first axis of the PCA on the distribution of the other species. “^2^” designates the quadratic effect of a variable. A star on top of the estimate indicates significance (p < 0.05); a dot indicates a trend (0.10 > p > 0.05).

## 4 DISCUSSION

Designing efficient conservation measures for endangered species requires a good knowledge of their distribution and ecology. Yet, for inconspicuous species, traditional survey methods often fail to provide reliable distribution and ecological data. Here, we studied the fine-scale distribution and breeding habitat preferences of amphibians in a continuous wetland expense by means of eDNA metabarcoding. We recovered DNA from six out of seven local amphibian species, i.e., four native species, *L. vulgaris, R. temporaria, B. bufo, H. arborea*, and two non-native species from the *Pelophylax* complex, *P. ridibundus* and *P. bergeri*. However, we did not detect DNA from one local species *L. helveticus*. We then found significant positive and negative associations between several environmental variables and some of the detected taxa, highlighting the potential of this approach to understand the ecology of a species in these natural systems, and thus provide data for reliable long-term monitoring.

In particular, our multivariate analysis showed that the main factor structuring species communities was water temperature. In addition, the proportion of emerged land and emerged vegetation both had a marginal impact on communities. Preserving microhabitat diversity thus appears key to maintaining community diversity.

We then investigated the specific habitat characteristics of each of our focal species. The fact that *L. helveticus* was not detected in areas where *L. vulgaris* is present suggests strong habitat segregation between these related species. This contrasts with previous findings of co-occurrence and similar niches at other locations (Griffiths, 1986, 1987). This could be explained by *L. vulgaris* being an opportunistic species which can occupy a wide range of habitats when available (Ćirović et al., 2008; Dolmen, 1988). Accordingly, it was only marginally more likely to be found closer to the wintering habitat, in line with previous findings that it does not migrate over large distances (Kovar et al., 2009), and other environmental variables did not seem to influence its distribution. Interestingly, studies elsewhere suggested that vegetation cover might be associated with the presence of *L. vulgaris*. Some studies found that a high coverage in vegetation is required for *L. vulgaris* to breed (Babik & Rafiński, 2001; Cooke & Frazer, 1976; Ildos & Ancona, 1994), while Marnell, (1998) found that the presence of *L. vulgaris* was associated with open water bodies with less floating vegetation, although he acknowledged that submerged grasses were more common in ponds where this species was present. Other environmental parameters such as the chemical composition of the water might affect habitat use by these species. Indeed, Cooke & Frazer (1976) suggested that *L. vulgaris* and *L. helveticus* tend to occupy water bodies with different chemical properties. While the latter is associated with soft and acidic water, *L. vulgaris* might occur more in hard and neutral water. Taken together, these findings suggest that *L. vulgaris* is a generalist species that can use different breeding habitat types and migrate for up to three hundred meters from the forest (Table S1), while *L. helveticus* would be restricted to wetlands located closer to the wintering habitats.

The only factor influencing the distribution of *R. temporaria* was the distance to the wintering habitats. *R. temporaria* is a generalist species that can breed in a wide range of temporal and permanent water bodies (Babik & Rafiński, 2001; Beebee, 1981; Marnell, 1998), in agreement with the results from our eDNA approach. In addition, this species breeds early in the season, and some froglets might already have emancipated from the area where they were born at the time of sampling.

The distribution of *P. ridibundus* was influenced by mud depth. This is consistent with the fact that most *Pelophylax* species often overwinter in the mud (Kovar et al., 2009; Tattersall & Ultsch, 2008). Consequently, the low densities measured during the prenuptial migration survey might be underestimated, as a large proportion of individuals might have overwintered downstream from the barriers.

Interestingly, the distributions of *P. ridibundus* and the related *P. bergeri* were not influenced by the same environmental factors. *P. bergeri* was found in areas with high vegetation cover and tended to be found in areas occupied by fewer other anurans. Whether this reflects competitive exclusion or niche segregation is unclear. This invasive species has replaced the native *P. lessonae* in our study area and throughout a large part of its range (Dufresnes et al., 2017), but its impact on other species is unknown. Importantly, *P. ridibundus* can produce F1 hybrids with *P. lessonae* or *P. bergeri*. Because we targeted a mitochondrial marker, it was not possible to identify hybrids. Consequently, some individuals labeled as *P. ridibundus* or *P. bergeri* might be hybrids, which might have increased heterogeneity in habitat characteristics within these two species, thus decreasing our ability to detect significant associations.

We did not detect a significant association between the presence of either *B. bufo* or *H. arborea* and any of the environmental variables recorded. However, *H. arborea* tended to be found mainly in habitats with lower emerged vegetation. Higher statistical power is required to confirm this trend, as *H. arborea* was found at only four sampling points.

We did not find a positive correlation between the RRA and the estimated amphibian species densities, contrary to what has been demonstrated repeatedly (Di Muri et al., 2020; Handley et al., 2019; Hänfling et al., 2016; Li et al., 2021b). However, as pointed out by Di Muri et al. (2020), standard survey methods have their own biases. Indeed, the migration barriers used in this study were not installed continuously during the migration period, which could have led to an underestimation of species densities. In addition, this survey method is ineffective for *P. ridibundus*, which overwinters primarily underwater, as well as for *H. arborea*, which is able to climb the barriers. The use of eDNA thus appears as a good complement to traditional migration surveys to assess more fine-scale distribution.

Three main hypotheses could explain the absence of *L. helveticus* in our water samples. First, it could be absent from the two reserves we sampled. This species is indeed known to be absent from the wetlands of Gletterens, however it is expected to be present in Yverdon where we detected 97 individuals during the prenuptial migratory survey. Second, the density of *L. helveticus* individuals in Yverdon might be too low to enable its detection using eDNA approaches. This would be unlikely, as we detected other local amphibian species from which fewer individuals were recorded during the prenuptial migration (Table S1). In addition, it has been repeatedly demonstrated that eDNA is a sensitive method for detecting organisms at low densities, as is the case for the detection of alien species in the initial phase of invasion (Cowart et al., 2018; Dejean et al., 2012; Jerde et al., 2011). Finally, the absence of *L. helveticus* DNA in our water samples could be the result of this species breeding near the wintering habitats, in places not covered by our sample points. In this study, we targeted three main vegetation types that are considered most suitable for breeding for most amphibian species. However, other temporarily flooded vegetation types such as *Nanocyperion, Cladietum* and *Caricion davallianae* (Delarze et al., 1998) are found directly downstream of the migration barriers and could be used as breeding habitats by *L. helveticus*. This hypothesis is in line with the literature, as this species has been previously observed to winter near its breeding habitat (Diego-Rasilla & Luengo, 2007).

In contrast, we detected *H. arborea* DNA in two sampling points in Yverdon. This result is surprising, as this species has not been observed in this natural reserve since 2000 (data from the Swiss Centre for Wildlife Mapping; https://lepus.unine.ch/carto/70120) and no individual was captured during the prenuptial migration survey at this location. This occurrence is hard to explain by cross-contaminations, as none of the samples were collected or extracted at the same time as the samples from Gletterens containing *H. arborea* DNA. Cross-contaminations during the PCR amplification also seem unlikely, as the samples where this species was detected were spread throughout the plate layout. It is thus likely that DNA from this species was indeed present in Yverdon. It is possible, though unlikely, that this species is still present at a low density in the area but has been undetected for decades. Alternatively, the detected DNA could originate from older eDNA resuspended from the sediments by other animals.

The application of eDNA-based approaches to study fine-scale habitat use in a continuous wet meadow expanse poses new challenges due to the biological and environmental conditions that influence eDNA dynamics. For instance, DNA persistence must be considered to determine the temporal and spatial scales at which environmental variables must be measured to describe the species habitat. Previous studies have investigated the persistence of DNA in water in laboratory or mesocosm conditions and established that DNA remains detectable approximately up to two weeks in the water column (Thomsen et al., 2012), while its persistence can extend over several years in sediments (Corinaldesi et al., 2008) or soil (Foucher et al., 2020). In the present study, we aimed to study habitat use of our focal amphibian species during reproduction. Thus, to avoid contamination from older DNA, we recorded the environmental variables one month before the collection of water samples to ensure DNA resuspended from the sediments had sufficient time to degrade, and we were careful not to resuspend the sediments during sampling. Another crucial parameter to design eDNA surveys is the diffusion potential of DNA in the environment caused by hydrodynamics processes affecting water currents. Brys et al. (2021) determined that the distance to detect amphibians using eDNA methods ranges from five meters for a newt to up to ten meters for frogs at high density. In *La Grande Cariçaie*, the water bodies are interspaced with stretches of emerged land as well as dense vegetation, which reduces DNA diffusion in the environment. We therefore considered that the five meter radius we used to characterize the habitat was a realistic estimate.

Our study highlighted the potential of an eDNA-based approach to investigate species ecology at a fine scale in a continuous wet meadow expanse. Overall, our results showed that eDNA metabarcoding allows the study of species’ habitat choice, leading to a better understanding of their ecology. Increased knowledge of species ecology is crucial for designing effective conservation policies to protect endangered species by conserving and restoring threatened environments.

## Acknowledgments

We thank all members of the Association of the Grande Cariçaie, particularly Alexandre Ghiraldi for his help in setting up field work and Christian Clerc for sharing the vegetation map of the reserves. We thank Pierre Taberlet for his advice on eDNA extraction, amplification, and sequencing methods; Nadège Remollino and Céline Stoffel for their help and advices during the lab work; Eduard Mas Carrio for his advices on bioinformatic and statistical analyses; Judith Schneider for her feedback on the results. This study was funded by the Association de la Grande Cariçaie, Cheseaux-Noréaz, and the School of Biology, University of Lausanne.

## Data and code availability

The DNA metabarcoding data generated for this study is available on SRA (BioProject ID: PRJNA938759, availability upon acceptance of the manuscript). Sanger sequences for *L. helveticus* and *P. bergeri* generated for this study were deposited in GenBank under accession nr. OK032503 and OQ508898, respectively. All data, OBITools script and R script underlying the study can be downloaded from the public repository: https://github.com/JulieGuenat/Amphibian_habitat_Characterization_eDNA.

## Authors contribution

(i) Conception and design of the study: JMG, AG, GL, LF; (ii) Acquisition, analysis, and interpretation of the data: JMG & GL; (iii) First draft JMG & GL; and (iv) Manuscript Editing JMG, AG, LF & GL.

## References

Arntzen, J. W., Abrahams, C., Meilink, W. R. M., Iosif, R., & Zuiderwijk, A. (2017). Amphibian decline, pond loss and reduced population connectivity under agricultural intensification over a 38 year period. Biodiversity, 26(6), 1411–1430. https://doi.org/10.1007/s10531-017-1307-y

Babik, W., & Rafiński, J. (2001). Amphibian breeding site characteristics in the western Carpathians, Poland. Herpetological Journal, 11(2), 41–51.

Barnes, M. A., & Turner, C. R. (2016). The ecology of environmental DNA and implications for conservation genetics. Conservation Genetics, 17, 1–17. https://doi.org/10.1007/s10592-015-0775-4

Bates, D., Mächler, M., Bolker, B. M., & Walker, S. C. (2015). Fitting linear mixed-effects models using lme4. Journal of Statistical Software, 67(1). https://doi.org/10.18637/jss.v067.i01

Becker, C. G., Fonseca, C. R., Haddad, C. F. B., & Prado, P. I. (2010). Habitat split as a cause of local population declines of amphibians with aquatic larvae. Conservation Biology, 24(1), 287–294. https://doi.org/10.1111/j.1523-1739.2009.01324.x

Beebee, T. J. C. (1981). Habitats of the British amphibians (4): Agricultural lowlands and a general discussion of requirements. Biological Conservation, 21(2), 127–139. https://doi.org/10.1016/0006-3207(81)90075-6

Bellemain, E., Carlsen, T., Brochmann, C., Coissac, E., Taberlet, P., & Kauserud, H. (2010). ITS as an environmental DNA barcode for fungi: An in silico approach reveals potential PCR biases. BMC Microbiology, 10(1), 189. https://doi.org/10.1186/1471-2180-10-189

Biggs, J., Ewald, N., Valentini, A., Gaboriaud, C., Dejean, T., Griffiths, R. A., Foster, J., Wilkinson, J. W., Arnell, A., Brotherton, P., Williams, P., & Dunn, F. (2015). Using eDNA to develop a national citizen science-based monitoring programme for the great crested newt (Triturus cristatus). Biological Conservation, 183, 19–28. https://doi.org/10.1016/j.biocon.2014.11.029

Boyer, F., Mercier, C., Bonin, A., Le Bras, Y., Taberlet, P., & Coissac, E. (2016). OBITOOLS: A unix-inspired software package for DNA metabarcoding. Molecular Ecology Resources, 16(1), 176–182. https://doi.org/10.1111/1755-0998.12428

Brys, R., Haegeman, A., Halfmaerten, D., Neyrinck, S., Staelens, A., Auwerx, J., & Ruttink, T. (2021). Monitoring of spatiotemporal occupancy patterns of fish and amphibian species in a lentic aquatic system using environmental DNA. Molecular Ecology, 30(13), 3097–3110. https://doi.org/10.1111/mec.15742

Calderón-Sanou, I., Münkemüller, T., Zinger, L., Schimann, H., Yoccoz, N. G., Gielly, L., Foulquier, A., Hedde, M., Ohlmann, M., Roy, M., Si-Moussi, S., & Thuiller, W. (2021). Cascading effects of moth outbreaks on subarctic soil food webs. Scientific Reports, 11(1). https://doi.org/10.1038/s41598-021-94227-z

Chalmandrier, L., Pansu, J., Zinger, L., Boyer, F., Coissac, E., Génin, A., Gielly, L., Lavergne, S., Legay, N., Schilling, V., Taberlet, P., Münkemüller, T., & Thuiller, W. (2019). Environmental and biotic drivers of soil microbial β-diversity across spatial and phylogenetic scales. Ecography, 42(12), 2144–2156. https://doi.org/10.1111/ECOG.04492

Ćirovi ć, R., Radovi ć, D., & Vukov, T. D. (2008). Breeding site traits of European newts (Triturus macedonicus, Lissotriton vulgaris, and Mesotriton alpestris: Salamandridae) in the Montenegrin karst region. Archives of Biological Sciences, 60(3), 459–468. https://doi.org/10.2298/ABS0803459C

Cooke, A. S., & Frazer, J. F. D. (1976). Characteristics of newt breeding sites. Journal of Zoology, 178(2), 223–236.

Corinaldesi, C., Beolchini, F., & Dell’Anno, A. (2008). Damage and degradation rates of extracellular DNA in marine sediments: Implications for the preservation of gene sequences. Molecular Ecology, 17(17), 3939–3951. https://doi.org/10.1111/j.1365-294X.2008.03880.x

Cowart, D. A., Breedveld, K. G. H., Ellis, M. J., Hull, J. M., & Larson, E. R. (2018). Environmental DNA (eDNA) applications for the conservation of imperiled crayfish (Decapoda: Astacidea) through monitoring of invasive species barriers and relocated populations. Journal of Crustacean Biology, 38(3), 257–266. https://doi.org/10.1093/jcbiol/ruy007

Cristescu, B., & Boyce, M. S. (2013). Focusing ecological research for conservation. In Ambio (Vol. 42, Issue 7, pp. 805–815). Springer. https://doi.org/10.1007/s13280-013-0410-x

Davidson, N. C. (2014). How much wetland has the world lost? Long-term and recent trends in global wetland area. Marine and Freshwater Research, 65(10), 934–941. https://doi.org/10.1071/MF14173

De Barba, M., Miquel, C., Boyer, F., Mercier, C., Rioux, D., Coissac, E., & Taberlet, P. (2014). DNA metabarcoding multiplexing and validation of data accuracy for diet assessment: Application to omnivorous diet. Molecular Ecology Resources, 14(2), 306–323. https://doi.org/10.1111/1755-0998.12188

Dejean, T., Valentini, A., Duparc, A., Pellier-Cuit, S., Pompanon, F., Taberlet, P., & Miaud, C. (2011). Persistence of environmental DNA in freshwater ecosystems. PLoS ONE, 6(8), e23398. https://doi.org/10.1371/journal.pone.0023398

Dejean, T., Valentini, A., Miquel, C., Taberlet, P., Bellemain, E., & Miaud, C. (2012). Improved detection of an alien invasive species through environmental DNA barcoding: The example of the American bullfrog Lithobates catesbeianus. Journal of Applied Ecology, 49(4), 953–959. https://doi.org/10.1111/j.1365-2664.2012.02171.x

Delarze, R., Gonseth, Y., & Galland, P. (1998). Guide des milieux naturels de Suisse: écologie, menaces, espèces caractéristiques (Delachaux et niestlé (ed.)).

Di Muri, C., Handley, L. L., Bean, C. W., Li, J., Peirson, G., Sellers, G. S., Walsh, K., Watson, H. V., Winfield, I. J., & Hänfling, B. (2020). Read counts from environmental DNA (eDNA) metabarcoding reflect fish abundance and biomass in drained ponds. Metabarcoding and Metagenomics, 4, 97–112. https://doi.org/10.3897/MBMG.4.56959

Diego-Rasilla, F. J., & Luengo, R. M. (2007). Acoustic orientation in the palmate newt, Lissotriton helveticus. Behavioral Ecology and Sociobiology, 61(9), 1329–1335. https://doi.org/10.1007/s00265-007-0363-9

Dolmen, D. (1988). Coexistence and niche segregation in the newts Triturus vulgaris (L.) and T. cristatus (Laurenti). Amphibia-Reptilia, 9(4), 365–374.

Dubey, S., Leuenberger, J., & Perrin, N. (2014). Multiple origins of invasive and “native” water frogs (Pelophylax spp.) in Switzerland. Biological Journal of the Linnean Society, 112(3), 442–449. https://doi.org/10.1111/bij.12283

Dufresnes, C., Déjean, T., Zumbach, S., Schmidt, B. R., Fumagalli, L., Ramseier, P., & Dubey, S. (2019). Early detection and spatial monitoring of an emerging biological invasion by population genetics and environmental DNA metabarcoding. Conservation Science and Practice, 1(9). https://doi.org/10.1111/csp2.86

Dufresnes, C., Di Santo, L., Leuenberger, J., Schuerch, J., Mazepa, G., Grandjean, N., Canestrelli, D., Perrin, N., & Dubey, S. (2017). Cryptic invasion of Italian pool frogs (Pelophylax bergeri) across Western Europe unraveled by multilocus phylogeography. Biological Invasions, 19(5), 1407–1420. https://doi.org/10.1007/s10530-016-1359-z

Eiler, A., Löfgren, A., Hjerne, O., Nordén, S., & Saetre, P. (2018). Environmental DNA (eDNA) detects the pool frog (Pelophylax lessonae) at times when traditional monitoring methods are insensitive. Scientific Reports, 8(1), 1–9. https://doi.org/10.1038/s41598-018-23740-5

Ficetola, G. F., Coissac, E., Zundel, S., Riaz, T., Shehzad, W., Bessière, J., Taberlet, P., & Pompanon, F. (2010). An In silico approach for the evaluation of DNA barcodes. BMC Genomics, 11(1), 434. https://doi.org/10.1186/1471-2164-11-434

Ficetola, G. F., Manenti, R., & Taberlet, P. (2019). Environmental DNA and metabarcoding for the study of amphibians and reptiles: species distribution, the microbiome, and much more. In Amphibia Reptilia (Vol. 40, Issue 2, pp. 129–148). Brill. https://doi.org/10.1163/15685381-20191194

Ficetola, G. F., Miaud, C., Pompanon, F., & Taberlet, P. (2008). Species detection using environmental DNA from water samples. Biology Letters, 4(4), 423–425. https://doi.org/10.1098/rsbl.2008.0118

Foucher, A., Evrard, O., Ficetola, G. F., Gielly, L., Poulain, J., Giguet-Covex, C., Laceby, J. P., Salvador-Blanes, S., Cerdan, O., & Poulenard, J. (2020). Persistence of environmental DNA in cultivated soils: implication of this memory effect for reconstructing the dynamics of land use and cover changes. Scientific Reports, 10(1), 1–12. https://doi.org/10.1038/s41598-020-67452-1

Garlapati, D., Charankumar, B., Ramu, K., Madeswaran, P., & Ramana Murthy, M. V. (2019). A review on the applications and recent advances in environmental DNA (eDNA) metagenomics. In Reviews in Environmental Science and Biotechnology (Vol. 18, Issue 3, pp. 389–411). https://doi.org/10.1007/s11157-019-09501-4

Goldberg, C. S., Turner, C. R., Deiner, K., Klymus, K. E., Thomsen, P. F., Murphy, M. A., Spear, S. F., McKee, A., Oyler-McCance, S. J., Cornman, R. S., Laramie, M. B., Mahon, A. R., Lance, R. F., Pilliod, D. S., Strickler, K. M., Waits, L. P., Fremier, A. K., Takahara, T., Herder, J. E., & Taberlet, P. (2016). Critical considerations for the application of environmental DNA methods to detect aquatic species. In M. Gilbert (Ed.), Methods in Ecology and Evolution (Vol. 7, Issue 11, pp. 1299–1307). John Wiley & Sons, Ltd (10.1111). https://doi.org/10.1111/2041-210X.12595

Griffiths, R. A. (1986). Feeding Niche Overlap and Food Selection in Smooth and Palmate Newts, Triturus vulgaris and T. helveticus, at a Pond in Mid-Wales. Journal of Animal Ecology, 55(1), 201–214. https://doi.org/10.2307/4702

Griffiths, R. A. (1987). Microhabitat selection and feeding relations of smooth and warty newts, Triturus vulgaris and T. cristatus, at an upland pond in mid-Wales. Journal of Animal Ecology, 56(2), 441–151. https://doi.org/10.1111/j.1600-0587.1987.tb00731.x

Handley, L. L., Read, D. S., Winfield, I. J., Kimbell, H., Johnson, H., Li, J., Hahn, C., Blackman, R., Wilcox, R., Donnelly, R., Szitenberg, A., & Hänfling, B. (2019). Temporal and spatial variation in distribution of fish environmental DNA in England’s largest lake. Environmental DNA, 1(1), 26–39. https://doi.org/10.1002/edn3.5

Hänfling, B., Handley, L. L., Read, D. S., Hahn, C., Li, J., Nichols, P., Blackman, R. C., Oliver, A., & Winfield, I. J. (2016). Environmental DNA metabarcoding of lake fish communities reflects long-term data from established survey methods. Molecular Ecology, 25(13), 3101–3119. https://doi.org/10.1111/mec.13660

Harper, L. R., Lawson Handley, L., Hahn, C., Boonham, N., Rees, H. C., Gough, K. C., Lewis, E., Adams, I. P., Brotherton, P., Phillips, S., & Hänfling, B. (2018). Needle in a haystack? A comparison of eDNA metabarcoding and targeted qPCR for detection of the great crested newt (Triturus cristatus). Ecology and Evolution, 8(12), 6330–6341. https://doi.org/10.1002/ece3.4013

Hartig, F. (2020). DHARMa: Residual Diagnostics for Hierarchical (Multi-Level / Mixed) Regression Models. http://florianhartig.github.io/DHARMa/

Houlahan, J. E., Fidlay, C. S., Schmidt, B. R., Meyer, A. H., & Kuzmin, S. L. (2000). Quantitative evidence for global amphibian population declines. Nature, 404(6779), 752–755. https://doi.org/10.1038/35008052

Ildos, A. S., & Ancona, N. (1994). Analysis of amphibian habitat preferences in a farmland area (Po plain, northern Italy). Amphibia Reptilia, 15(3), 307–316. https://doi.org/10.1163/156853894X00083

Jerde, C. L., Mahon, A. R., Chadderton, W. L., & Lodge, D. M. (2011). “Sight-unseen” detection of rare aquatic species using environmental DNA. Conservation Letters, 4(2), 150–157. https://doi.org/10.1111/j.1755-263X.2010.00158.x

Kovar, R., Brabec, M., Bocek, R., & Vita, R. (2009). Spring migration distances of some Central European amphibian species. Amphibia Reptilia, 30(3), 367–378. https://doi.org/10.1163/156853809788795236

Legendre, P., & Andersson, M. J. (1999). Distance-based redundancy analysis: Testing multispecies responses in multifactorial ecological experiments. Ecological Monographs, 69(1), 1–24. https://doi.org/10.1890/0012-9615(1999)069[0001:DBRATM]2.0.CO;2

Li, W., Hou, X., Xu, C., Qin, M., Wang, S., Wei, L., Wang, Y., Liu, X., & Li, Y. (2021b). Validating eDNA measurements of the richness and abundance of anurans at a large scale. Journal of Animal Ecology, 90(6), 1466–1479. https://doi.org/10.1111/1365-2656.13468

Li, W., Song, T., Hou, X., Qin, M., Xu, C., & Li, Y. (2021a). Application of eDNA metabarcoding for detecting anura on a tropical island. Diversity, 13(9), 440. https://doi.org/10.3390/d13090440

Lodge, D. M., Turner, C. R., Jerde, C. L., Barnes, M. A., Chadderton, L., Egan, S. P., Feder, J. L., Mahon, A. R., & Pfrender, M. E. (2012). Conservation in a cup of water: Estimating biodiversity and population abundance from environmental DNA. Molecular Ecology, 21(11), 2555–2558. https://doi.org/10.1111/j.1365-294X.2012.05600.x

Lopes, C. M., Sasso, T., Valentini, A., Dejean, T., Martins, M., Zamudio, K. R., & Haddad, C. F. B. (2017). eDNA metabarcoding: a promising method for anuran surveys in highly diverse tropical forests. Molecular Ecology Resources, 17(5), 904–914. https://doi.org/10.1111/1755-0998.12643

Marnell, F. (1998). Discriminant analysis of the terrestrial and aquatic habitat determinants of the smooth newt (Triturus vulgaris) and the common frog (Rana temporaria) in Ireland. Journal of Zoology, 244(1), 1–6. https://doi.org/10.1111/j.1469-7998.1998.tb00001.x

Martinez-Almoyna, C., Thuiller, W., Chalmandrier, L., Ohlmann, M., Foulquier, A., Clément, J. C., Zinger, L., & Münkemüller, T. (2019). Multi-trophic β-diversity mediates the effect of environmental gradients on the turnover of multiple ecosystem functions. Functional Ecology, 33(10), 2053–2064. https://doi.org/10.1111/1365-2435.13393

Mas-Carrió, E., Schneider, J., Nasanbat, B., Ravchig, S., Buxton, M., Nyamukondiwa, C., Stoffel, C., Augugliaro, C., Ceacero, F., Taberlet, P., Glaizot, O., Christe, P., & Fumagalli, L. (2022). Assessing environmental DNA metabarcoding and camera trap surveys as complementary tools for biomonitoring of remote desert water bodies. Environmental DNA, 4(3), 580–595. https://doi.org/10.1002/edn3.274

Meyer, A., Zumbach, S., Schmidt, B., & Monney, J.-C. (2009). Les amphibiens et les reptiles de Suisse. Haupt.

Nielsen, K. M., Johnsen, P. J., Bensasson, D., & Daffonchio, D. (2007). Release and persistence of extracellular DNA in the environment. In Environmental Biosafety Research (Vol. 6, Issues 1–2, pp. 37–53). EDP Sciences. https://doi.org/10.1051/ebr:2007031

Oksanen, J., Simpson, G., Blanchet, F., Kindt, R., Legendre, P., Minchin, P., O’Hara, R., Solymos, P., Stevens, M., Szoecs, E., Wagner, H., Barbour, M., Bedward, M., Bolker, B., Borcard, D., Carvalho, G., Chirico, M., De Cacares, M., Durand, S., … Weedon, J. (2020). vegan: Community Ecology Package (R package version 2.5-7).

Perl, R. G. B., Avidor, E., Roll, U., Malka, Y., Geffen, E., & Gafny, S. (2022). Using eDNA presence/non-detection data to characterize the abiotic and biotic habitat requirements of a rare, elusive amphibian. Environmental DNA, 4(3), 642–653. https://doi.org/10.1002/edn3.276

Pilliod, D. S., Goldberg, C. S., Arkle, R. S., & Waits, L. P. (2013). Estimating occupancy and abundance of stream amphibians using environmental DNA from filtered water samples. Canadian Journal of Fisheries and Aquatic Sciences, 70(8), 1123–1130. https://doi.org/10.1139/cjfas-2013-0047

Pilliod, D. S., Goldberg, C. S., Arkle, R. S., & Waits, L. P. (2014). Factors influencing detection of eDNA from a stream-dwelling amphibian. Molecular Ecology Resources, 14(1), 109–116. https://doi.org/10.1111/1755-0998.12159

R core Team, & R Foundation for Statistical Computing. (2020). R: A Language and Environment for Statistical Computing (3.4.4). https://www.r-project.org/

Rödel, M.-O., & Ernst, R. (2004). Measuring and monitoring amphibian diversity in tropical forests. I. An evaluation of methods with recommendations for standardization. Ecotropica, 10(1), 1–14.

Ruppert, K. M., Kline, R. J., & Rahman, M. S. (2019). Past, present, and future perspectives of environmental DNA (eDNA) metabarcoding: A systematic review in methods, monitoring, and applications of global eDNA. In Global Ecology and Conservation (Vol. 17, p. e00547). https://doi.org/10.1016/j.gecco.2019.e00547

Sakata, M. K., Maki, N., Sugiyama, H., & Minamoto, T. (2017). Identifying a breeding habitat of a critically endangered fish, Acheilognathus typus, in a natural river in Japan. Science of Nature, 104(11–12), 1–8. https://doi.org/10.1007/S00114-017-1521-1

Shehzad, W., Riaz, T., Nawaz, M. A., Miquel, C., Poillot, C., Shah, S. A., Pompanon, F., Coissac, E., & Taberlet, P. (2012). Carnivore diet analysis based on next-generation sequencing: application to the leopard cat (Prionailurus bengalensis) in Pakistan. Molecular Ecology, 21(8), 1951–1965. https://doi.org/10.1111/j.1365-294X.2011.05424.x

Smart, A. S., Tingley, R., Weeks, A. R., van Rooyen, A. R., & McCarthy, M. A. (2015). Environmental DNA sampling is more sensitive than a traditional survey technique for detecting an aquatic invader. Ecological Applications, 25(7), 1944–1952. https://doi.org/10.1890/14-1751.1

Taberlet, P., Bonin, A., Zinger, L., & Coissac, E. (2018). Environmental DNA: For biodiversity research and monitoring. Oxford University Press. https://doi.org/10.1093/oso/9780198767220.001.0001

Taberlet, P., Coissac, E., Hajibabaei, M., & Rieseberg, L. H. (2012). Environmental DNA. Molecular Ecology, 21(8), 1789–1793. https://doi.org/10.1111/j.1365-294X.2012.05542.x

Tattersall, G. J., & Ultsch, G. R. (2008). Physiological Ecology of Aquatic Overwintering in Ranid Frogs. In Biological Reviews (Vol. 83, Issue 2, pp. 119–140). John Wiley & Sons, Ltd. https://doi.org/10.1111/j.1469-185X.2008.00035.x

Thomsen, P. F., Kielgast, J., Iversen, L. L., Wiuf, C., Rasmussen, M., Gilbert, M. T. P., Orlando, L., & Willerslev, E. (2012). Monitoring endangered freshwater biodiversity using environmental DNA. Molecular Ecology, 21(11), 2565–2573. https://doi.org/10.1111/j.1365-294X.2011.05418.x

Valentini, A., Taberlet, P., Miaud, C., Civade, R., Herder, J., Thomsen, P. F., Bellemain, E., Besnard, A., Coissac, E., Boyer, F., Gaboriaud, C., Jean, P., Poulet, N., Roset, N., Copp, G. H., Geniez, P., Pont, D., Argillier, C., Baudoin, J. M., … Dejean, T. (2016). Next-generation monitoring of aquatic biodiversity using environmental DNA metabarcoding. Molecular Ecology, 25(4), 929–942. https://doi.org/10.1111/mec.13428

Wang, C., Qian, L., Zhang, C., Guo, W., Pan, T., Wu, J., Wang, H., & Zhang, B. (2017). A new species of Rana from the Dabie Mountains in eastern China (Anura, Ranidae). ZooKeys, 724, 135–153. https://doi.org/10.3897/zookeys.724.19383

Willerslev, E., Hansen, A. J., Binladen, J., Brand, T. B., Gilbert, M. T. P., Shapiro, B., Bunce, M., Wiuf, C., Gilichinsky, D. A., & Cooper, A. (2003). Diverse plant and animal genetic records from holocene and pleistocene sediments. Science, 300(5620), 791–795. https://doi.org/10.1126/science.1084114

Zedan, H. (2004). 2004 IUCN red list of threatened species: a global species assessment (J. Baillie, C. Hilton-Taylor, & S. N. Stuart (eds.)). IUCN--The World Conservation Union. https://doi.org/10.2305/iucn.ch.2005.3.en

