## Supplementary material for "Investigating fine-scale breeding habitat use by amphibians in a continuous wetland using environmental DNA"

By Julie M. Guenat et al.

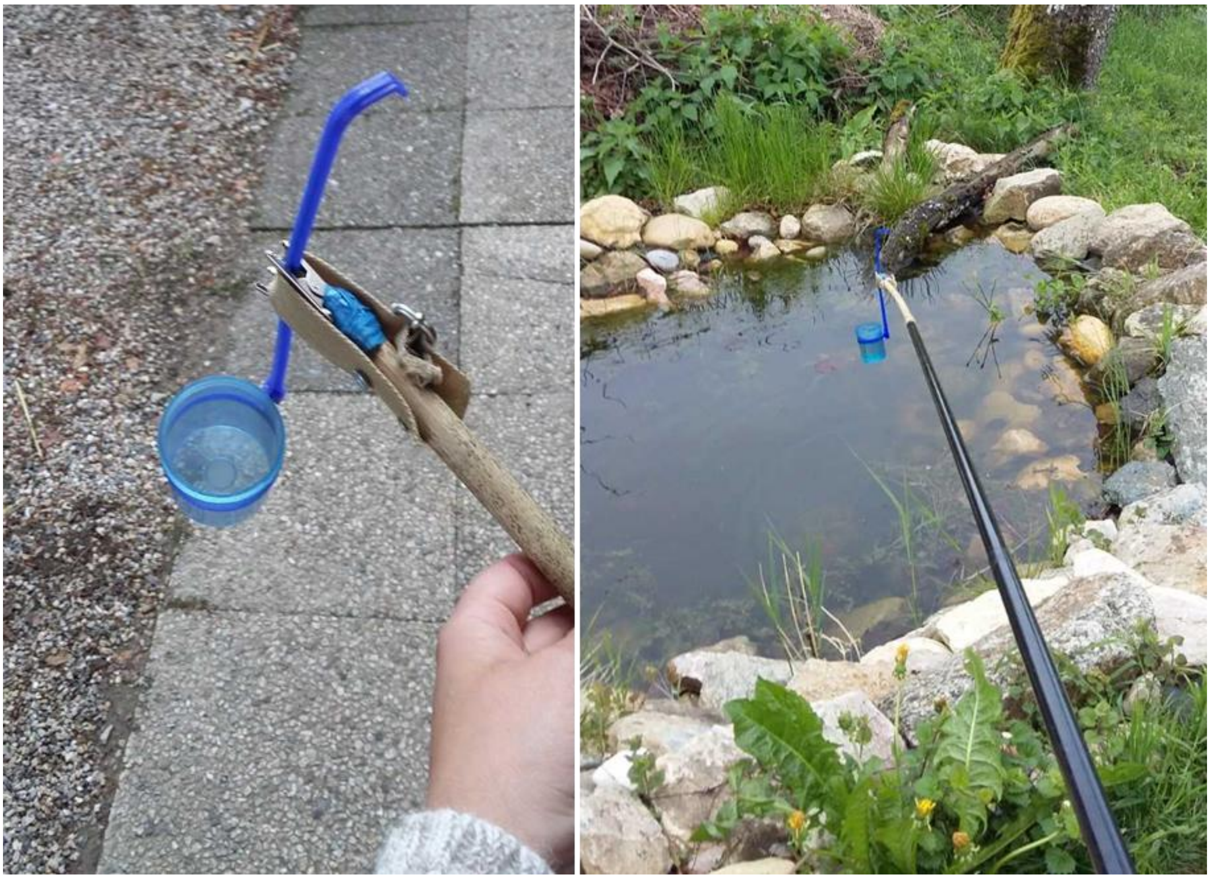

**Fig. S1: Sampling material.** Spoon from the kit VigiDNA (Spygen) attached to the four-meter fishing rod by means of two electric grippers and a belt. The water body presented on the right picture does not reflect environmental conditions of the Grande Cariçaie wetlands. Pictures were taken in Fontanezier (Switzerland, VD)

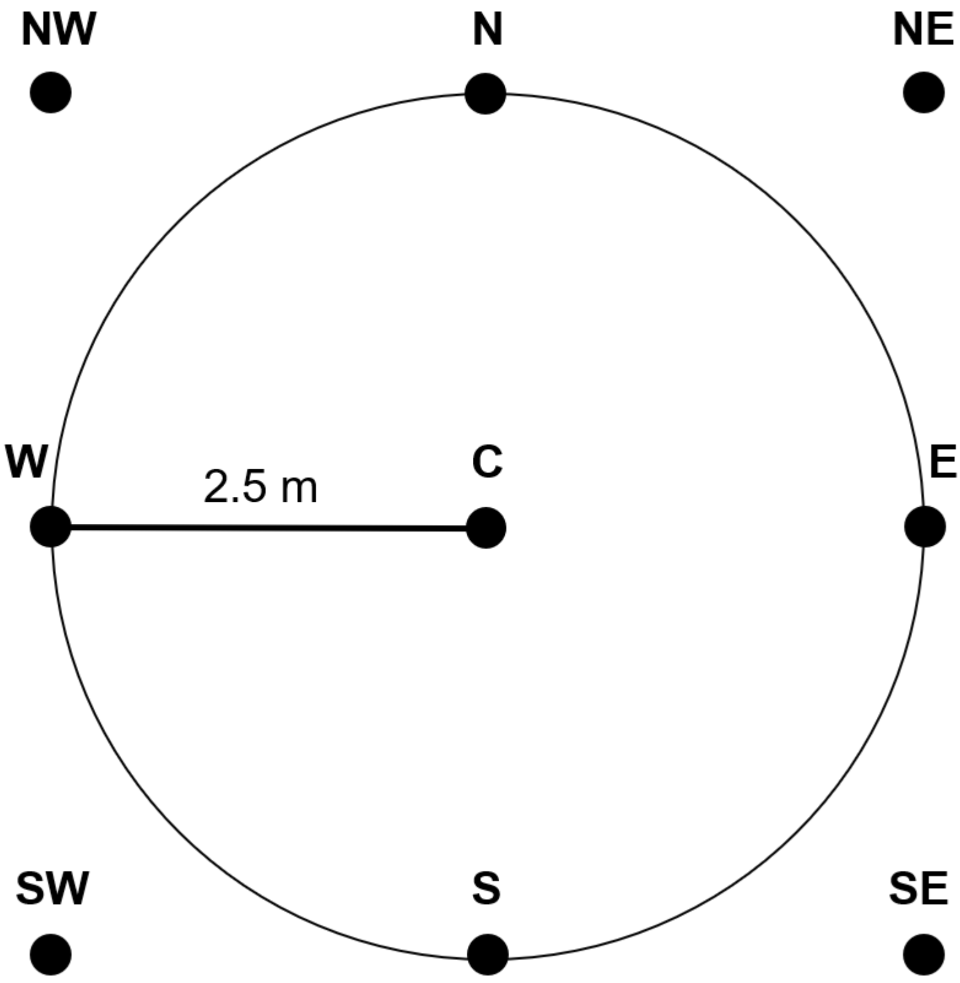


**Fig. S2: Scheme of water and mud depth measurements strategy performed at each sampling point.** Measurements were taken at 2.5 m from the center. C = Center; W = West; NW = Northwest; N = North; NE = Northeast; E = East; SE = Southeast; S = South; SW = Southwest. The circle represents the sampling point with a diameter of 5 m.

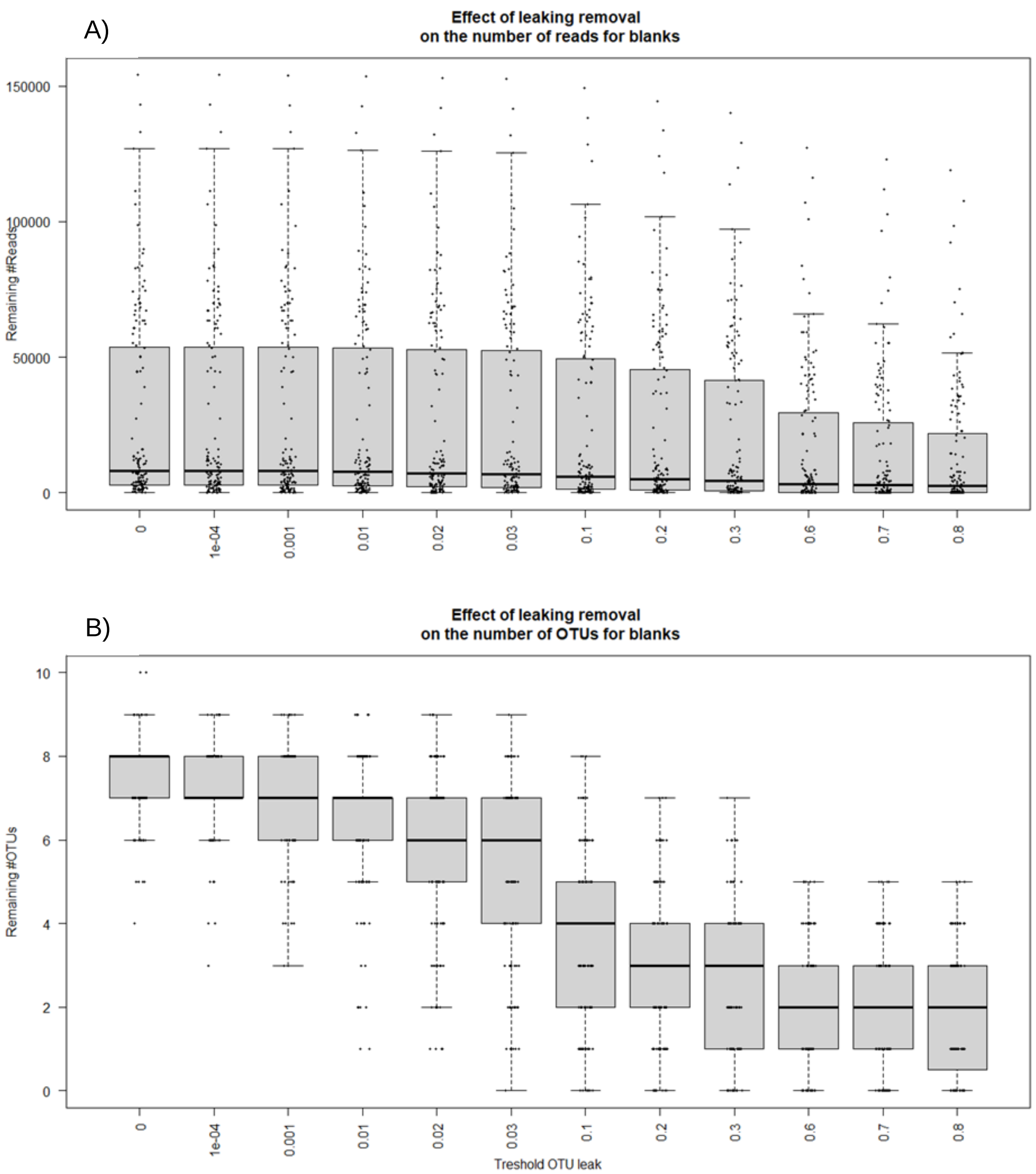


**Fig. S3: Effect of Leaking removal on the number of A) Reads B) OTUs in Blanks.** The x axis corresponds at proportion of A) sequences and B) OTUs removal. The y axis corresponds at the number of remaining A) sequences and B) OTUs in blanks. We observe on both graph that removing more than 60% of A) sequences and B) OTUs does not reduce the number of remaining A) sequences and B) OTUs in blanks. Thus, we removed 60% of total reads per BATR01 MOTUs in each sample to account for contaminants caused by sequence leaking.

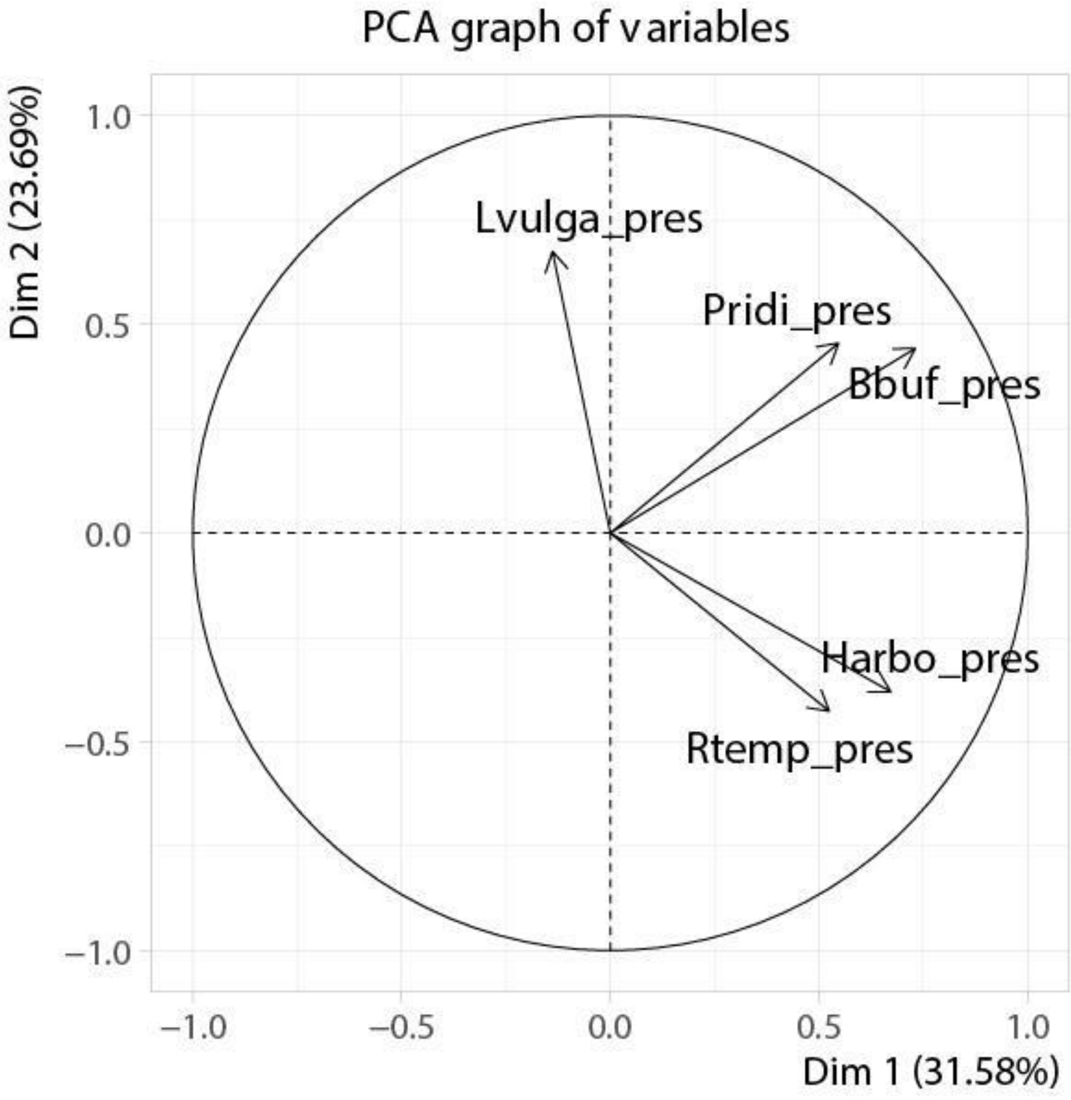


**Fig. S4: Contribution of each species to community structure at points where *P. bergeri* was found, excluding this species.** Vectors represent the contribution of each species to overall community structure. PC1 was used as a variable in GLMs (see main text).

**Table S1: Summary of the number of amphibian individuals detected with different methods**. N is the number of individuals and n is the number of sampling points. The surface of the sampling area in Yverdon is 12.3 ha and in Gletterens 17.3 ha; thus the density of sampling points per hectare is 2.03 in Yverdon and 1.44 in Gletterens. The density has not been estimated for *P. ridibundus* as this species mostly does not overwinter upstream from our migration barriers.

| ***Species*** | **Individuals captured during the prenuptial migration survey** | | | | **Number of sampling points where the species was detected (out of 25)** | | | **Maximum distance to wintering habitats [m]** |
|---|---|---|---|---|---|---|---|---|
| | **Yverdon** | | **Gletterens** | | **Yverdon** | **Gletterens** | **Total** | |
| | **N** | **Density [N * ha$^{-1}$]** | **N** | **Density [N * ha$^{-1}$]** | **n** | **n** | | |
| *L. vulgaris* | 443 | 36.02 | 97 | 5.61 | 5 | 7 | 12 | 296.97 |
| *L. helveticus* | 97 | 7.89 | 0 | 0 | - | - | - | - |
| *B. bufo* | 55 | 4.47 | 131 | 7.57 | 9 | 10 | 19 | 376.89 |
| *P. ridibundus* | 0.3 | 0.02 | 0.4 | 0.02 | 3 | 3 | 6 | 376.89 |
| *P. bergeri* | 0.7 | 0.06 | 1.6 | 0.09 | 10 | 7 | 17 | 390.60 |
| *R. temporaria* | 139 | 11.3 | 76 | 4.39 | 8 | 20 | 28 | 390.60 |
| *H. arborea* | 0 | 0 | 9 | 0.52 | 2 | 2 | 4 | 390.60 |

**Table S2: Results of the permutational multivariate analysis of variance (permanova) performed on distance based-Redundancy Analysis (db-RDA).**

| Variables | DF | Sum of squares | $R^2$ | F | Pr (>F) |
|---|---|---|---|---|---|
| **Submerged Vegetation** | 1 | 0.0711 | 0.00505 | 0.2600 | 0.869 |
| **(Submerged Vegetation)$^2$** | 1 | 0.0207 | 0.00147 | 0.0756 | 0.923 |
| **Emerged Vegetation** | 1 | 0.7685 | 0.05456 | 2.8091 | *0.060* |
| **(Emerged Vegetation)$^2$** | 1 | 0.5378 | 0.03818 | 1.9658 | 0.132 |
| **Emerged Land** | 1 | 0.2706 | 0.01921 | 0.9892 | 0.397 |
| **(Emerged Land)$^2$** | 1 | 0.6429 | 0.04565 | 2.3500 | *0.084* |
| **Mud Depth** | 1 | 0.1046 | 0.00743 | 0.3825 | 0.763 |
| **(Mud Depth)$^2$** | 1 | 0.2242 | 0.01592 | 0.8194 | 0.522 |
| **Forest Distance** | 1 | 0.3352 | 0.02380 | 1.2251 | 0.318 |
| **(Forest Distance)$^2$** | 1 | 0.1656 | 0.01175 | 0.6052 | 0.643 |
| **Water temperature** | 1 | 0.0662 | 0.00470 | 0.2421 | 0.818 |
| **(Water temperature)$^2$** | 1 | 0.7548 | 0.05359 | 2.7588 | ***0.045*** |
| **Residuals** | 37 | 10.1228 | 0.71869 | | |
| **Total** | 49 | 14.0851 | 1.00000 | | |

### Supplementary methods S1: Comparison of temperature measurements using 1-Wire®/iButton or Onset Hobo® thermologgers

Water temperature was measured every hour at each sampling point from May 1$^{st}$ to July 1$^{st}$, 2018 using thermologgers (1-Wire®/iButton®). Since these thermologgers are not waterproof, they were placed in Falcon tubes sealed with parafilm (hereafter Falcon thermologgers). To investigate the potential bias induced by Falcon tubes, waterproof thermologgers (Onset Hobo®) were also placed at three representative sampling points to get the direct water temperature. For two of the three sampling points, the temperature records between waterproof and Falcon thermologgers did not differ. A larger variation in temperature records was observed for the third sampling point (Fig. SM1.1). This sampling point was assumed to be an outlier and temperature records from the waterproof and Falcon thermologgers were assumed to be generally equal. Hence, for other sampling points, temperature from Falcon thermologgers were taken as such.

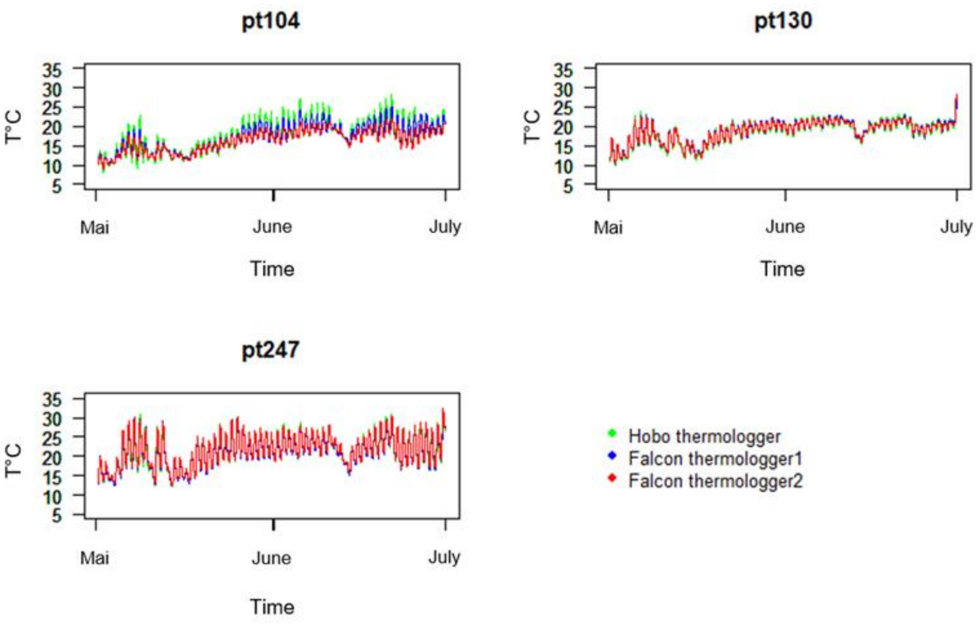


**Fig. SM1.1: Comparison of temperature records between the waterproof thermologger (Onset Hobo®) and the two thermologgers (1-Wire®/iButton®) contained in Falcon tubes at three sampling points.** Temperature was recorded from May $1^{st}$ to July $1^{st}$. Pt104 was in the *Magnocaricion* in the Yverdon reserve. Pt130 was in the *Phragmition* in the Yverdon reserve. Pt247 was in the *Nymphaeion* in the Gletterens reserve.

**Supplementary methods S2: Positive control**

We added a positive control to seven wells of each of the 12 PCR plates during the amplification of the 12S mitochondrial gene of amphibians using BATR01 primers.

We aimed at obtaining similar DNA concentration of the positive control to that of the eDNA samples. For this purpose, we compared the CT results obtained during the RT-qPCR we performed to test sample inhibitions of the dilution curve to those of the environmental samples (Table SM2.1). The dilution curve consisted in a serial dilution from $10^{-0}$x to $10^{-6}$x of *Rana iberica* DNA extracted from tissues (initial concentration Table SM2.2). From Table SM2.1 we observed that the mean CT obtained for the environmental samples is closer to the CT obtained with dilution $10^{-5}$x than that obtained with the dilution $10^{-6}$x of the dilution curve. Hence, we decided to adjust the concentration in the positive control to the $10^{-5}$x diluted *R. iberica* concentration.

The positive control consisted in an equimolar assembly of DNA from three exotic species, namely *Pelophylax nigromaculatus*, *Pseudacris maculata* and *Rana arvalis*. DNA from these species were extracted from tissues and these samples were quantified using Qubit® 2.0 Fluorometer (Life Technology Corporation; Table SM2.2).

We needed a total of 300 µL at a DNA concentration of 47.2 x $10^{-5}$ ng/µL of positive control. Thus, it represented per species: 100 µL (Final Volume) at a concentration of 47.2 x 1/33333 ng/µL (Final Concentration). To compute the volume of DNA solution (Initial volume, Vi) of each of the three species composing the positive control we used the following equation where Ci correspond at the initial DNA concentration quantified using Qbit:

$$Vi = \frac{47.2 * \frac{1}{33333} * 100}{Ci}$$

Results for each species are shown in Table SM2.2.

**Table SM2.1: Mean CT obtained for the dilution curve and the eight tested samples during the RT-QPCR used to test environmental samples for inhibition.**

| Dilution curve –Mean CT | | Samples –Mean CT |
|---|---|---|
| **Dilution** | **CT** | |
| $10^{-0}$x | 17.129 | 36.499 |
| $10^{-1}$x | 20.497 | |
| $10^{-2}$x | 24.134 | |
| $10^{-3}$x | 27.896 | |
| $10^{-4}$x | 31.864 | |
| $10^{-5}$x | 34.796 | |
| $10^{-6}$x | 39.330 | |
| $H_2O$ | 41.334 | |

**Table SM2.2: Results of the quantification of *P. nigromaculatus, P. maculata* and *R. arvalis* using Qubit® 2.0 Fluorometer.**

| Species | Concentration [ng/µL] | Volume required [µL] |
|---|---|---|
| *R. iberica* $10^{-1}$ x | 47.2 | |
| *R. iberica* $10^{-5}$ x | 47.2 x $10^{-5}$ | |
| *P. nigromaculatus* $10^{-1}$ x | 2.84 | 5 x $10^{-2}$ |
| *P. maculata* $10^{-1}$ x | 10.10 | 14 x $10^{-3}$ |
| *R. arvalis* $10^{-1}$ x | 27.30 | 5.2 x $10^{-3}$ |

**Supplementary methods S3: *Lissotriton helveticus* 12S partial gene sequenced using Sanger sequencing.**

Since the interest portion of the 12 S mitochondrial gene of *L. helveticus* was missing in EMBL (fig. SM3.1), we sequenced it using Sanger sequencing. We performed a PCR on extracted *L. helveticus* tissues. The PCR mixture contained 1 U of AmpliTaq Gold polymerase, 1x PCR gold buffer, 2 mM of $MgCl_2$, 0.2 mM of each dNTPs, 0.5 µM of forward and reverse primers, 0.2 mg/mL of bovine serum albumin and 2 µL of template DNA, resulting in a final volume of 25 µL. Thermocycling conditions were as follows; denaturation and activation of the polymerase at 95 °C for 10 min, followed by 30 cycles of 30 s at 95 °C, 30 s at 50 °C and 1 min at 72 °C, followed by a final elongation at 72 °C for 7 min.

We performed then a nested PCR to ensure that amplicons contain the targeted amplicon from amplification with BATR01. Same PCR mixture was used and the thermocycling conditions were 10 min at 95°C for DNA denaturation, followed by 10 cycles of 30 s at 95°C, 30 s at 55°C and 1 min at 72°C, followed by a final elongation of 7 min at 72°C.

Amplicon amplified with primers L2519 and H3296 was then sequenced using Sanger sequencing.

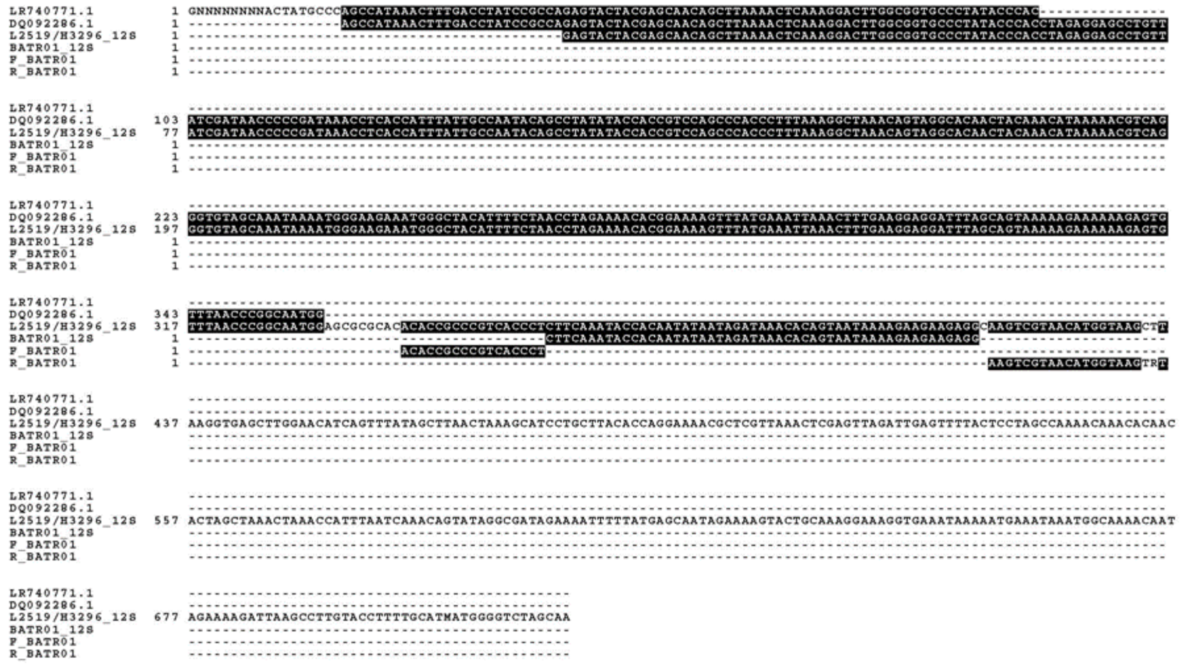

**Fig. SM3.1: Alignment of 12S mitochondrial sequences of *L. helveticus*.**
LR740771.1 and DQ092286.1 correspond to the *L. helveticus* mitochondrial 12S gene sequences extracted from GenBank. L2519/H3296_12S corresponds to the 12S mitochondrial gene sequence we amplified using primers L2519/H3296 from Wang et al. 2017. BATR01_12S corresponds to the 12S mitochondrial gene sequence that is amplified using BATR01, thus represents the target amplicon. F_BATR01 and R_BATR01 represent respectively the forward and the reverse BATR01 primer from Valentini et al. 2016.

### Supplementary methods S4: Investigating the absence of *Lissotriton helveticus* DNA in water samples.

While the absence of *L. helveticus* was expected in Gletterens, we were assuming to detect DNA of this species in water samples from Yverdon.

Thus, we tested to map sequences from water samples to the sequenced 12S mitochondrial partial gene of *L. helveticus* using *bwa* and *samtools*. 36 sequences were found to match the *L. helveticus* 12S partial gene (hereafter matching sequences). To investigate phylogenetical distances among the 36 matching sequences and the *L. helveticus* 12S partial gene, a tree was constructed using MEGA (figure SM4.1). The 12S mitochondrial partial gene of *L. helveticus* was found to be an outgroup of matching sequences. The 36 matching sequences are shown to be grouped with the *L. vulgaris* 12S partial gene.

**Fig. SM4.1: Phylogenetical distances between 12S and 16S partial gene of L.**

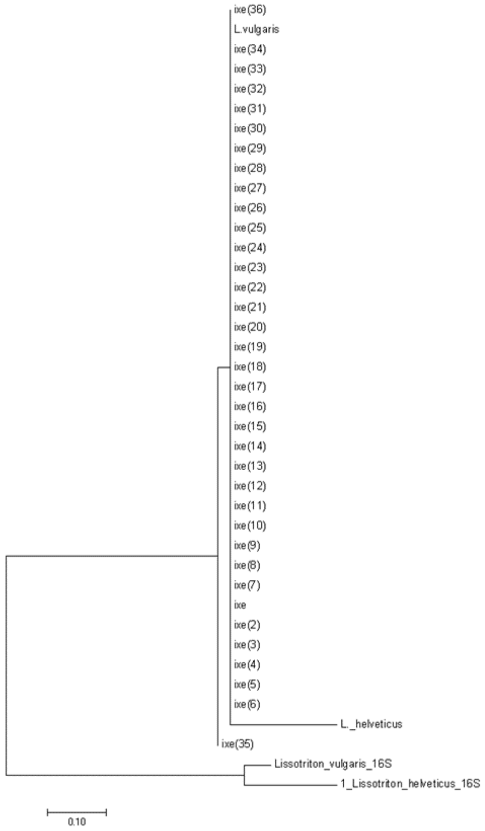


**helveticus and L. vulgaris as well as the 36 matching sequences recovered using**

**samtools**. Ixe(1-36) correspond to matching sequences. L_helveticus corresponds to the 12s partial sequence of this species. L_vulgaris corresponds to the 12S partial gene sequence of this species.

### Supplementary information S1: Investigation of presences of *H. arborea* in Yverdon

DNA from *H. arborea* was detected in the Yverdon reserves, while no individuals were captured during the migration survey. According to the data from the Swiss Centre for Wildlife Mapping (CSCF, https://lepus.unine.ch/carto/70120), the last observation of this species was in 2000 in this reserve. These presences could be attributed to cross-contamination during the handling of the samples in the field and in the laboratory.

However, this is unlikely as none of the Gletterens water samples were collected or extracted at the same time as Yverdon water samples containing *H. arborea* DNA (Table SI1.1). This excludes a cross-contamination between samples from Gletterens and those from yverdon.

During the amplification of samples, one sample out of the five positives to *H. arborea* from Yverdon was surrounded by samples from Gletterens, excluding the hypothesis of a sequence leaking during this manipulation (Fig. SI1.1).

**Table SI1.1: Date of water collection, DNA precipitation, extraction, and extraction series number of the nine samples where DNA of *H. arborea* was detected.** The number of reads retrieved for each sample after filtering and binding PCR replicates are shown.

| Location | Sample ID | Water collection | DNA precipitation | Extraction | Extraction series# |
|---|---|---|---|---|---|
| Yverdon | 107 | 25.05.18 | 23.08.18 | 27.08.18 | 17 |
| Yverdon | 125 | 23.05.18 | 16.08.18 | 17.08.18 | 3 |
| Gletterens | 217 | 27.05.18 | 22.08.18 | 24.08.18 | 12 |
| Gletterens | 242 | 24.05.18 | 20.08.18 | 24.08.18 | 11 |

**Plan de plaques PCR Metabarcoding** JG

| | 1 | 2 | 3 | 4 | 5 | 6 | 7 | 8 | 9 | 10 | 11 | 12 |
|---|---|---|---|---|---|---|---|---|---|---|---|---|
| A | B1 | E01 | 200 | 205 | N03 | 225 S | 248 | P04 | 145 | 123 S | 108 | E07 |
| B | Font | B2 | 102 | 109 | P02 | 141 | 215 S | 133 | 211 | 215 | 100 | 223 |
| C | 125 | N01 | B3 | 201 | 203 | E04 | 249 | 252 S | N06 | 142 | 251 | P07 |
| D | 110 | P01 | 261 | B4 | 249 S | N04 | 129 | 130 | P05 | 224 | 111 | B12 |
| E | 207 | 217 | E02 | 127 S | B5 | P03 | 240 | 247 | 237 | E06 | B11 | 218 S |
| F | 231 | 242 | N02 | 252 | 124 S | B6 | E05 | 127 | 123 | B10 | 216 | 104 |
| G | 148 | 124 | 146 | 107 | 225 | 233 | B7 | 118 | B9 | N07 | 144 | 236 |
| H | 131 | 111 S | 140 | E03 | 218 | 118 S | N05 | B8 | 129 S | P06 | 147 | Cudrefin |

**Figure SI1.1**: **sample localization in the PCR plate.** White wells are samples, Grey wells are banks, Orange wells are extraction negative controls, Red wells are PCR negative control, Green wells are positive controls and Blue wells are samples for which *H. arborea* DNA was detected.
